# Antibiotic-induced DNA damage results in a controlled loss of pH homeostasis and genome instability

**DOI:** 10.1101/2020.01.27.921072

**Authors:** James Alexander Booth, Mário Špírek, Tekle Airgecho Lobie, Kirsten Skarstad, Lumir Krejci, Magnar Bjørås

**Author notes:** Corresponding Authors ( &). Contributed equally (Lumir Krejci, Magnar Bjørås).

## Abstract

Extracellular pH has been assumed to play little if any role in how bacteria respond to antibiotics and antibiotic resistance development. Here, we show that the intracellular pH of *Escherichia coli* equilibrates to the environmental pH following antibiotic treatment. We demonstrate that this allows the environmental pH to influence the transcription of various DNA damage response genes and physiological processes such as filamentation. Using purified RecA and a known pH-sensitive mutant variant RecA K250R we show how pH can affect the biochemical activity of a protein central to control of the bacterial DNA damage response system. Finally, two different mutagenesis assays indicate that environmental pH affects antibiotic resistance development. Specifically, at environmental pH’s greater than six we find that mutagenesis plays a significant role in producing antibiotic resistant mutants. At pH’s less than or equal to 6 the genome appears more stable but extensive filamentation is observed, a phenomenon that has previously been linked to increased survival in the presence of macrophages.

## INTRODUCTION

There is considerable medical concern about the rise in antibiotic-resistant infections (1). Recalcitrance to treatment has been shown to occur due to mutagenesis or mechanisms of transitory tolerance and persistence (2). Antibiotic treatment can lead to DNA damage, genomic instability and subsequently accelerated resistance development in bacteria. As a consequence of DNA damage the bacterial SOS response is induced (3, 4). The SOS response (SOS) encompasses over 50 genes with several linked to antibiotic resistance development (5, 6). SOS has several physiological effects including the depolarisation of the electrical potential difference (ΔΨ) across the inner membrane of the bacterium (7–9). ΔΨ quantifies the difference in charge across a membrane and is one of two components that make up the proton motive force (pmf). The proton gradient (ΔpH), the other component of the pmf varies as the external pH changes and pH homeostasis maintains the intracellular pH within a narrow range of ∼7.5-7.7 (10, 11). As bacteria such as *Escherichia coli* (*E. coli*) have a permissive proliferation pH range of 5-9 both ΔΨ and ΔpH have to be actively managed to maintain pmf and pH homeostasis (10).

Here, we wished to test if antibiotic treatment of *E. coli* would affect the ΔpH and to understand the consequences of these changes. First we demonstrated that the quinolone antibiotic, nalidixic acid, induces a *recA*-dependent controlled loss of ΔpH. As a consequence we show the changes in intracellular pH result in alterations in the biochemical activities of RecA, including recombination and strength of transcription of various SOS response genes. Additionally, filamentation, rates of mutagenesis and viability were also shown to be dependent on the external pH. Our results thus show that the pH of the external environment affects how the bacterium reacts to antibiotics and demonstrates that *E. coli* uses different antibiotic survival strategies at different pH’s.

## MATERIAL AND METHODS

### Strains and microbial techniques

Unless otherwise stated the wild type *E. coli* MG1655 strain was used. Bacterial strains and plasmids used are listed in Supplementary Table S1. The *recA* mutant was generated from the keio collection and transduced into MG1655 (12). Transfer of genotypes of interest was carried out by P1 transduction (13). Transformation of plasmids was carried out using electroporation using electrocompetent cells generated using a glycerol/mannitol density step centrifugation (14). Genetic antibiotic selection markers flanked by FRT sites were removed using the FLP recombinase containing plasmid pCP20 (15). Antibiotics were used at the following concentrations, kanamycin 50 µg/ml (plasmid selection) 30 µg/ml (genomic selection), ampicillin 100 µg/ml (plasmid selection) and nalidixic acid 100 µg/ml (optimum concentration for lethality) (16). LB broth (10 g tryptone, 5 g yeast extract, 10 g NaCl adjusted to 1 l with MQ H_2_O and autoclaved) was used for none pH adjusted experiments together with agar (1.5%) if for solid media. LBK (10 g tryptone, 5 g yeast extract, 7.45 g KCl, adjusted to 1 l with MQ H_2_O and autoclaved) was used to limit the sodium ion concentrations, which inhibit growth at high pH. M9 minimum glycerol media was prepared by diluting sterile (M9 (10x) 100 ml, MgSO_4_ (1 M) 1 ml, CaCl_2_ (0.2 M) 2 ml, thiamine (5%) 2 ml and glycerol (60%) 1.67 ml) into pre-autoclaved MQ H_2_O to 1 l, containing 15 g agar (1.5%) if for plates. M9 minimum lactose media was prepared as mentioned above by substituting lactose (20%, 50.1 ml) for glycerol. The pH’s of LBK, M9 minimal media and PBS solutions were adjusted (100 mM) with Good’s sulfonate buffers, 2-(N-Morpholino)ethanesulfonic acid hydrate (MES) for pH’s 5.2, 5.5 and 6, 3-(N-Morpholino)propanesulfonic acid (MOPS) for pH 7 and N-[Tris(hydroxymethyl)methyl]-3-aminopropanesulfonic acid (TAPS) for pH 8. Adjustments were carried out with KOH (5 M) and the solutions sterilised by filtration (0.2 µm).

### Survival assay – Nalidixic acid

Acute survival was measured by survival following the removal of the DNA damage induced by nalidixic acid. The strains were grown (37°C, 200 rpm) in quadruplet in media (LB or pH adjusted LBK, 3 ml in a 15 ml tube) to OD_600_=0.8. The first sample was taken before addition of nalidixic acid and designated as t=0. Samples were centrifuged (21,500 g, 20 s) media aspirated and the pellet re-suspended (PBS) to remove the nalidixic acid. The samples were then serially diluted (10x, 100 µl, PBS or pH adjusted PBS) in 96 well plates before samples (10 µl) were transferred to 24 well plates containing only LB-agar or pH adjusted LBK-agar (1 ml/well, 1.5% agar). The droplets were allowed to dry, then incubated (overnight, 37°C) in the dark before enumeration the following day. For each sample the well containing 10-100 colonies was counted and used to calculate CFU/ml. Samples were taken at the time points indicated on the graphs.

### Flow cytometry – Intracellualar pH determination

The strains were grown (37°C, 200 rpm) in quadruplet in pH adjusted media (LBK, 3 ml in a 15 ml tube) containing 200 µM arabinose to OD_600_=0.8 (Nanodrop, ND-1000). The arabinose maintains high levels of expression and keeps the majority of active GFP in the cytoplasm (17). The first sample was taken before addition of nalidixic acid and designated as t=0. The samples (2 µl) were diluted into several pH buffered PBS solutions (100 µl), one at the pH of the LBK and a calibration curve of five pH buffered PBS solutions containing the membrane permeable weak acid sodium benzoate (60 mM). Benzoate collapses the ΔpH across the inner membrane resulting in the internal pH equalization to that of the external solution. The extracellular pH’s were chosen to ensure that at least one of the solutions was higher and one was lower in pH compared to the intracellular pH of the bacteria. The fluorescence signal of the TorA-GFPmut3* protein from 10,000 bacteria/sample was then measured on an AccuriC6 flow cytometer in FLA-1. Subsequent samples were taken at the time points indicated after exposure to nalidixic acid. The intracellular pH values were calculated using the two solutions on the calibration curve that straddled the fluorescence intensity value of the unknown LBK sample.

### Flow cytometry - SOS induction and *recA, lexA & umuDC-gfp* measurement

The strains were grown (37°C, 200 rpm) in quadruplet in pH adjusted media (LBK, 3 ml in a 15 ml tube) to OD_600_=0.8 (Nanodrop, ND-1000). The first sample was taken before addition of nalidixic acid and designated as t=0. The samples (2 µl) were diluted into PBS (100 µl) buffered to the same pH as the incubating media and simultaneously at pH 8 supplemented with sodium benzoate (60 mM) at room temperature. The GFP signal is maximised and standardised by using PBS at pH 8 together with sodium benzoate (60 mM) and eliminates the GFP signal fluctuations due to the pH variations due to the pH equilibration. The fluorescence signal from 10,000 bacteria/sample from the GFPmut2 protein or the RecA-GFP fusion, where the GFP moiety is Gfp-901, a derivative of GFPmut2, were measured on an AccuriC6 flow cytometer in FLA-1. Subsequent samples were taken at the time points indicated after exposure to nalidixic acid.

### Electromobility shift assay (EMSA)

The EMSA described is a modification of a previously published protocol (18). RecA was diluted from concentrated stock (New England Biolabs M0249L) into Storage Buffer (50 mM phosphate/citrate buffer at pH 7, 50 mM NaCl), which was also used in no protein control. Protein was mixed with a master mix (containing 900 nM (nucleotides) 5’-FITC-labelled 90mer oligonucleotide (AAATCAATCTAAAGTATATATGAGTAAACTTGGTCTGACAGTTACCAATGCTTAATCAGTGAGGCACCTATCTCAGCGATCTGTCTATTT), 50 mM phosphate/citrate buffer at pH 5 to 8, 50 mM NaCl, 10 mM MgCl_2_ and 1 mM ATP) according to the indicated scheme in 10 μl reaction volume at 25°C for 5 minutes. The reactions were stopped by addition of glutaraldehyde to final concentration of 0.125% for additional 5 minutes and then transferred to ice. Reactions were resolved on 1% agarose gels in 1X TAE (70 V, 2 h). Gels were subjected to fluorescent analysis and data quantification using the FLA-9000 Starion (Fujifilm) and MultiGauge (Fujifilm) software.

### D-loop formation assays

The reaction was performed as described earlier (19). Briefly, RecA was incubated for 5 min with 50 nM (moles) 5’-FITC-labelled 90mer oligonucleotide as in the EMSA experiment and 2 μl (920 ng) of pBluescript SK(–) (460 ng/μl) was then added to bring the final reaction volume to 10 μl and incubated for 10 min at 37 °C. The samples were deproteinized with 0.1% SDS and 10 μg proteinase K for 10 min at 37°C and resolved in 0.9% agarose gels in 1X TAE (90 V, 35 min). Gels were imaged on a FLA-9000 scanner (Fujifilm) and quantified with Multi Gauge V3.2 (Fujifilm).

### Stopped-flow assays and data analysis

The experiments were performed essential as described previously using an SFM-300 stopped-flow machine (Bio-Logic) fitted with a MOS-200 monochromator spectrometer (Bio-Logic) with excitation wavelength set at 545 nm(20). Fluorescence measurements were collected with a 550 nm long pass emission filter. The machine temperature was maintained at 25°C with a circulating water bath. For all experimental setups, a master mix containing all common reaction components for each of the two syringes was prepared. To these mixtures a variable RecA amounts were added. Since equal volumes were injected into the mixing chamber from each syringe, the two solutions became mutually diluted. Therefore, all reaction components common to each syringe were prepared at the final concentration, whereas reaction components present in only one syringe were added at twice the desired final concentration. All concentrations quoted represent final concentrations after mixing. Components of each syringe were pre-incubated for 10 min before the start of experiments to allow the contents to reach equilibrium.

All reactions were performed in Stopped Flow Buffer (50 mM phosphate/citrate buffer (pH 5 to 8), 10 mM MgCl_2_, 50 mM NaCl, 1 mM ATP). All reactions contained 20 nM 5’-Cy3 fluorescently labelled (dT79). RecA was added directly from concentrated stocks. For all experiments, control reactions were also performed for buffer alone with and without DNA to confirm fluorescence signal stability over the time course of the experiments (data not shown).

Fluorescence measurements for most experiments were collected according to the following protocol: (1) every 0.00005 s from 0-0.05 s; (2) every 0.0005 s from 0.05-0.56 s; (3) every 0.02 s from 0.56-120.54 s. For each condition analysed, traces were collected from between three and nine independent reactions (n = 3-9) and averages were generated. Importantly, all conclusions from this study are made on the magnitude and rate of changes in fluorescence over time and how these vary with RecA concentration, and therefore the absolute fluorescence values were converted to arbitrary units by a normalisation procedure to facilitate comparison. For all experiments the raw data were normalised to the same fluorescence value for the 0 s time point.

For analysis, for all experiments ten-point moving averages were calculated on each individual normalised trace, which were used to define initial (0 s), final (120.54 s) and maximum fluorescence (and corresponding time point), and ΔCy3 fluorescence values for each experiment. To simplify direct comparison of conditions for various pH conditions we calculated observed overall half-times, measured from time points where the fluorescence from moving averages was closest to the value calculated for ΔCy3 fluorescence midpoints.

### LexA degradation assay

RecA (0.6 µM) was mixed with ssDNA (30-mer, 70 nM) in citrate-phosphate buffer with various pH containing 35 mM NaCl, 10 mM MgCl_2_ and either 1 mM ATP or ATPgammaS. RecA nucleofilaments were formed for 10 minutes at 37°C and then mixed with LexA protein. Reactions were stopped at 0, 60 or 120 minutes by addition of Laemmli buffer. Samples were resolved in 15% SDS-PAGE gels and stained with Coomassie stain. The intensity of bands for full length LexA protein were analysed by MultiGauge Software. The average signal is shown with standard error for 3 independent experiments.

### Lactose reversion assay

FC29 (Rif^S^, F’ Δ(*lacI lacZ*)) *E. coli* cannot utilise lactose as a carbon source or revert from Lac^-^ to Lac^+^. FC40 (Rif^R^, F’ *lacI*Ω*lacZ*) eliminates the coding sequence of the last four residues of *lacI*, all of *lacP* and *lacO*, and the first 23 residues of *lacZ* (21). Individual colonies on M9 minimum glycerol plates were used to seed M9 minimal glycerol (30°C, 200 rpm, 1 ml in 15 ml tubes, three days). pH adjusted M9 minimum lactose agar plates were inoculated with 10^10^ FC29 to scavenge each plate for sources of carbon other than lactose. The next day, the plates were overlaid with 1:10 (FC29:FC40) suspended in pH adjusted M9 minimal lactose top agar (0.5% agar) and incubated at 37°C.

The frequency of Lac^+^ revertants was determined by counting Lac^+^ colonies every 24 hours for 12 days and calculating the mean and standard deviation (n=3).

### MIC assay

Wild type *E. coli* (*MG1655*) was grown overnight (37°C, LB agar plates) before being resuspended in pH adjust LBK media and adjusted to an OD_600_=0.03. The antibiotics were dissolved to a final concentration of 128 µg/ml in pH adjusted LBK and serially diluted (x2) to 0.25 µg/ml (giving 10 concentrations of each antibiotic). A single µl of adjusted bacterial culture was added to each well (100 µl). The plates were then incubated (37°C) and the OD_595_ read the following day. The experiments were repeated in quadruplet to determine the mean and standard deviation.

### Forward UV induced rifampicin resistance assay

pH adjusted PBS and LBK media were used throughout. Wild type *E. coli* (*MG1655*) were grown overnight (37°C, 1.5% agar) before being diluted and resuspended (<1000 bacteria / culture) in liquid media for overnight growth (37°C, 200 rpm). The following day the bacteria were diluted (x50) and grown to OD_600_=0.25 before being centrifuged and resuspended in PBS and exposed to UV (20 J/m^2^) or not.

Subsequently all cultures were kept in the dark. The samples of the cultures were then plated on Rifampicin (100 µg/ml) plates and also on LBK to determine CFU. The cultures were then centrifuged and resuspended in LBK before incubation (37°C, 200 rpm, ON). The following day the cultures were plated on Rifampicin (100 µg/ml) and LBK to determine CFU. The following day the concentration of rifampicin resistant bacteria was calculated. The mean and standard deviation were calculated from three independent experiments.

### Bacterial viability assay with propidium iodide

Bacteria were grown overnight (37°C, LB agar plates) before being resuspended in pH adjust LBK media and adjusted to an OD_600_=0.03. The strain(s) were grown (37°C, 200 rpm) in pH adjusted media (LBK, 3 ml in a 15 ml tube) to OD_600_=0.6 (Nanodrop, ND-1000). The first sample was taken before addition of nalidixic acid and designated as t=0. The samples (2 µl) were diluted into PBS (100 µl) buffered to the sample pH as the incubating media and containing PI (2 µl) in pairs where one of the samples was also treated with EtOH (100 µl). The fluorescence signal of PI from 10,000 bacteria/sample were then measured on an AccuriC6 flow cytometer in FLA-3. Simultaneous readings of FSC-A were recorded and used to calculate the density of PI in the bacteria. Subsequent samples were taken at the time points indicated after exposure to nalidixic acid. All experiments were run on at least three independent days.

## RESULTS

### DNA damage leads to the loss of pH homeostasis

To induce DNA damage we used the antibiotic nalidixic acid a quinolone that binds DNA gyrase and topoisomerase IV inhibiting their ligation activities leading to DNA double strand breaks and chromosome fragmentation (22). To explore the hypothesis that antibiotic-induced DNA damage affects the proton gradient of the bacterium we first confirmed that nalidixic acid led to the *recA*-dependent loss of electrical potential (Supplementary Figure S1 and Results) (23). Thereafter, we tested the potential of a pH sensitive reporter, GFPmut3* (Supplementary Table S1), to follow intracellular pH changes (Supplementary Figure S2) (24). *E. coli* expressing GFPmut3* were grown to exponential phase before treating with Nalidixic acid (100 µg/ml). The fluorescence of GFPmut3* was recorded by flow cytometry both before and after treatment and simultaneously the GFPmut3* signal of each sample was calibrated over the possible pH range. We observed that the calibration curve changed over time following the introduction of the antibiotic and determined that continuous calibration of GFPmut3* was necessary (Supplementary Figure S3). The intracellular pH of *E. coli* was monitored for four hours following antibiotic-induced DNA damage where the external pH was buffered to pH 7 (Figure 1A). Following a delay of 30 minutes, the intracellular pH fell reaching the pH of the extracellular environment about 120 minutes after the addition of nalidixic acid.

**Figure 1.**
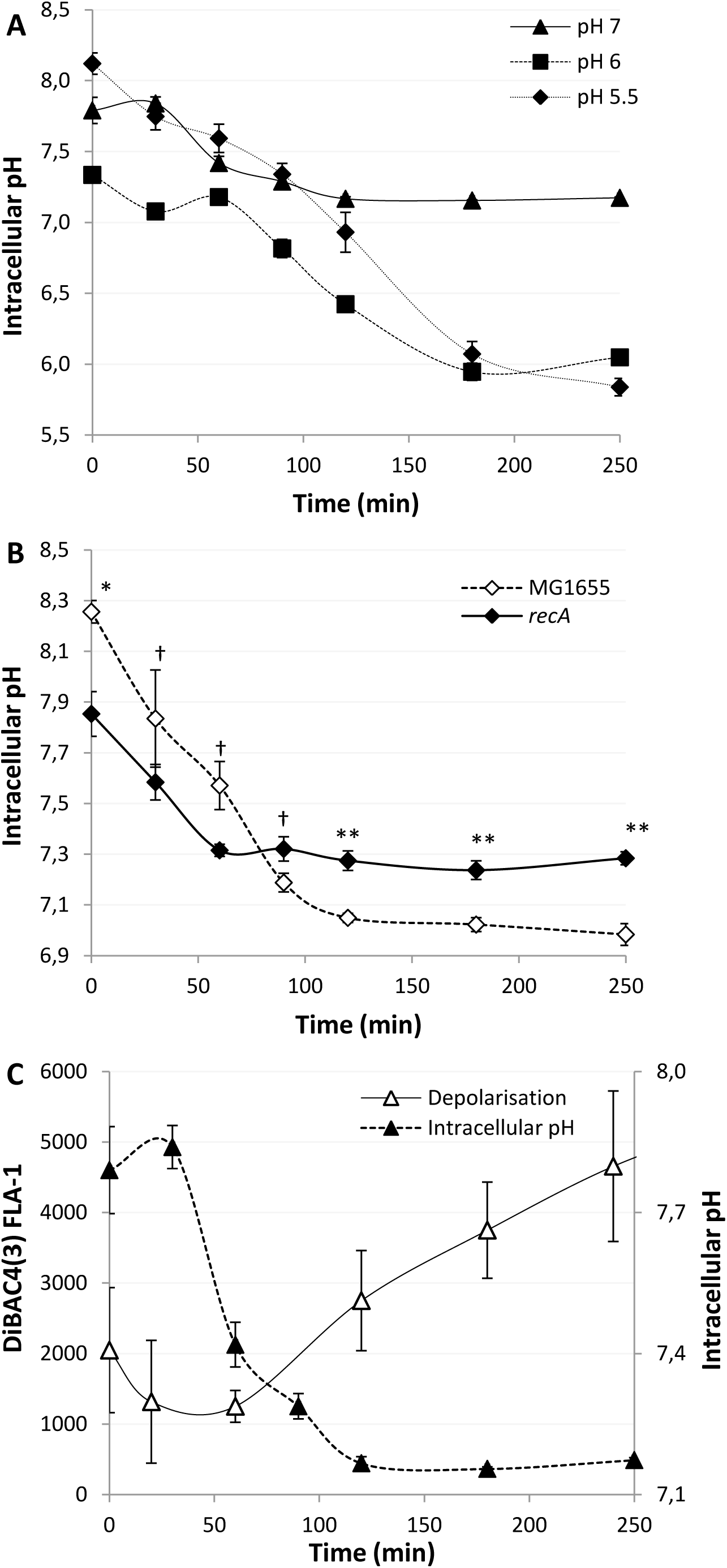
DNA damage leads to *recA* dependent loss of pH homeostasis. (**A-C**) The cytoplasmic pH of *E. coli* was measured using calibrated GFPmut3* fluorescent intensity measurements using flow cytometry following Nalidixic acid (100 µg/ml) induced DNA damage at 0 min. (**A**) pH depolarisation was followed over four hours, the extracellular media was buffered to 5.5, 6 and 7, as indicated on the graph. (**B**) pH equilibration is not complete in a *recA* mutant following exposure to Nalidixic acid at pH 7. (**C**) Intracellular pH equilibrates with the extracellular pH before electrical polarity as measured by DiBAC4(3) following DNA damage in wild type *E. coli*. All data points shown are mean ± s.e.m., *n* =4 (independent replicates). †p > 0.05, *p≤0.05, ** p ≤0.01 (t-test, two tail, unpaired equal variance).

*E. coli* is the predominant aerobic organism in the caecum and colon of the human gastrointestinal tract where the pH varies from 5.7-6.8 and 6.1-7.2 respectively (25, 26). Additionally, pathogenesis can lead certain *E. coli* strains to be exposed to the urinary tract (pH 4.5-8 (27)) and the eukaryotic intracellular environment which includes macrophages and phagosomes (pH 5-7.5 (28)). Due to these various ecological niches occupied by *E. coli*, we were curious as to how the external pH may affect the kinetics of loss of proton homeostasis in more acidic environments.

Buffering the media to either pH 6 or 5.5 led to the decrease of intracellular pH to almost that of the external media over 180-240 minutes (Figure 1A). As RecA coordinates the DNA damage induced-stress response in *E. coli*, by inducing the expression of LexA sensitive genes in a dose-dependent manner (29), we used a *recA* mutant to demonstrate a significant reduction in proton depolarisation after treatment with nalidixic acid as compared to wild type (Figure 1B). Additionally, we noted in the wild type that electrical depolarisation never fully equilibrated at pH 7 whilst proton equilibration was achieved after two hours. Identical experimental setups allowed for a direct comparison illustrating that the two processes appear to be kinetically independent (Figure 1C). Together these results show both ΔΨ and ΔpH equilibrate with their external environment in a *recA*-dependent manner. However, the kinetics of equilibration indicate the two processes appear be independent of each other.

### External pH affects filamentation and *recA* transcription

The *recA* dependency of proton equilibration following DNA damage suggested that various SOS processes may also be dependent on the external pH. Filamentation following induction of the SOS response is a classical morphological response to DNA damage (30–32). Therefore, we investigated if the external pH would influence the dynamics and extent of filamentation following DNA damage using flow cytometry by following forward scatter (FSC-A), as a proxy for cell volume(33). Surprisingly, when the extracellular pH was buffered to pH 7 after exposure to nalidixic acid cell filamentation showed an initial increase followed by a gradual decline (Figure 2A). Next we examined how the external pH buffered to 6 or 5.5 would affect the dynamics of filamentation. Both pH conditions showed a greater extent of filamentation (Figure 2A). These data demonstrate that following DNA damage the external pH can affect internal cellular processes, such as filamentation and loss of mean cell volume via changes in the intracellular pH.

**Figure 2.**
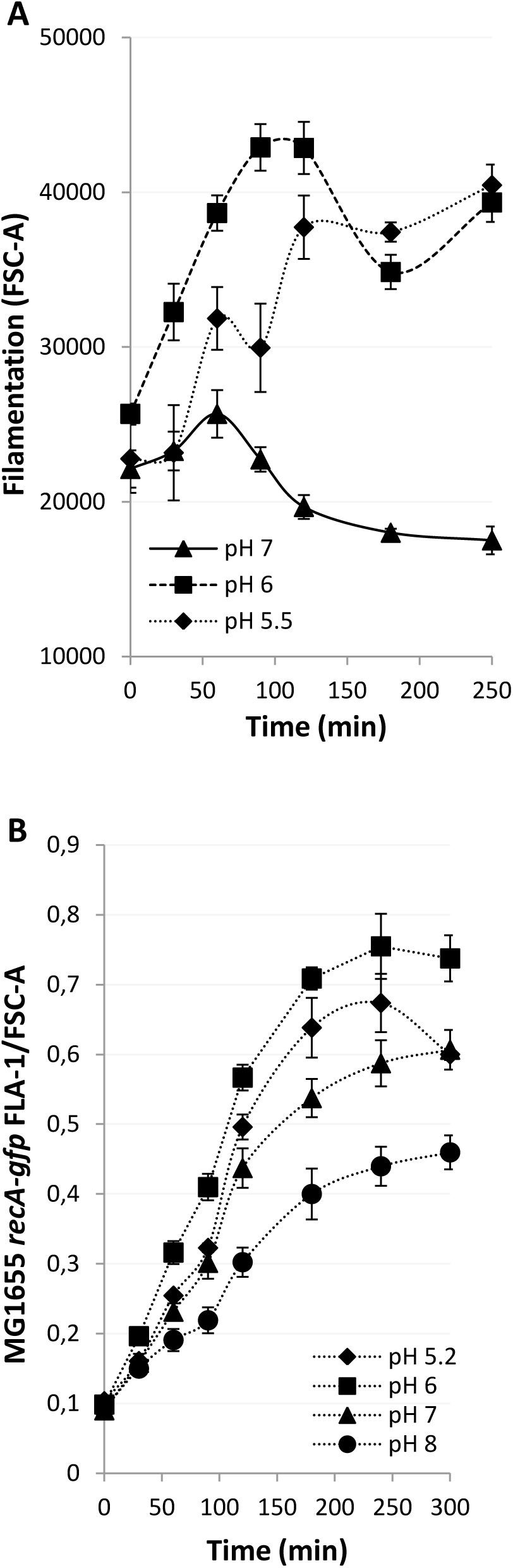
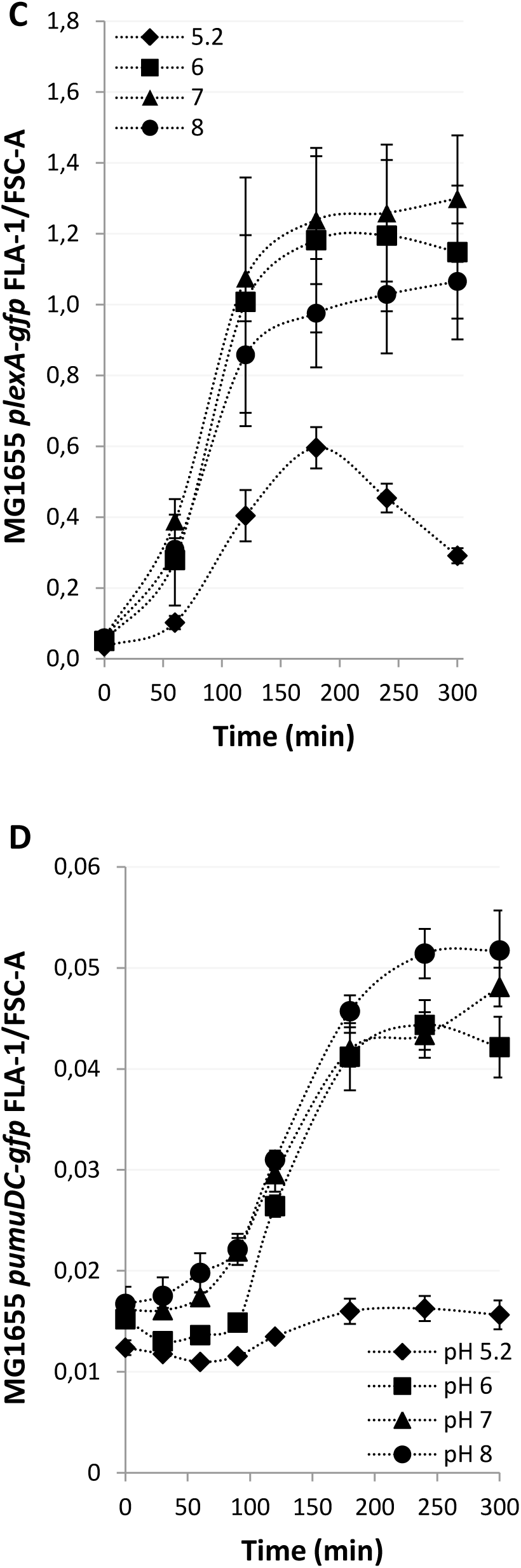
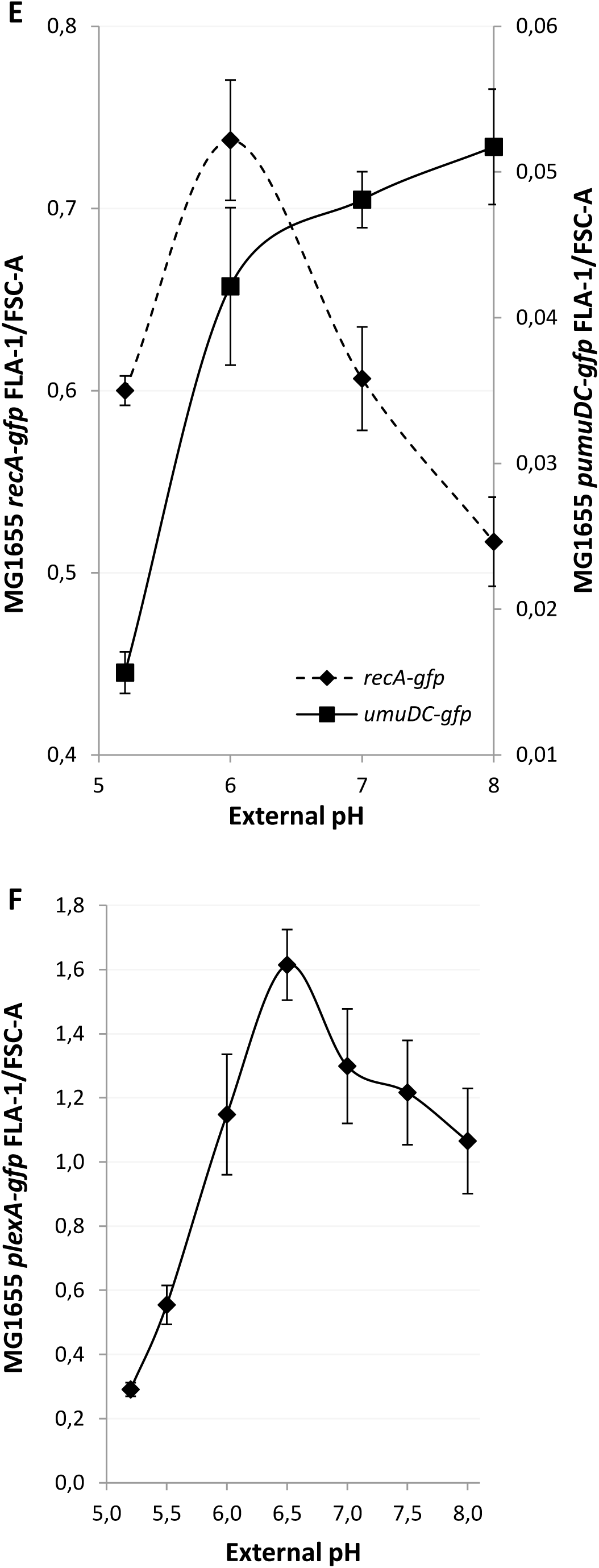
Proton depolarisation results in extracellular pH dependent affects on morphological plasticity and transcription of SOS genes *recA, lexA* and *umuDC*. (**A-D**) DNA damage was induced with Nalidixic acid (100 µg/ml) at 0 min and signals followed for 5 hours. (**A**) Kinetics and extent of filamentation, as defined by FSC-A, were followed by flow cytometry in wild type *E. coli*. (**B-D**) The liquid media was buffered to pH 5.2, 6, 7 and 8 and the gfpmut2 signal followed by flow cytometry. GFP fusions measured by fluorescence density measurements were (**B**) *recA-gfpmut2,* (**C**) *plexA - gfpmut2,* (**D**) *pumuDC -gfpmut2*. (**E**) The pH dependence of *recA* and the *umuDC* operon transcription following DNA damage (5 hours). (**F**) The pH dependence of *recA* transcription following DNA damage (5 hours). All data points are mean ± s.e.m., *n* =4 (independent replicates).

As the expression of RecA is immediate following the introduction of a substantial amount of DNA damage (34) we next examined the RecA-mediated transcriptional SOS response over the replicative permissive pH range of *E. coli*. To monitor the induction and kinetics of the SOS response we used a strain expressing an optimised genomic *recA-gfpmut2* fusion protein which responds quickly to SOS induction and minimizes steric interference of the tag (*recA-gfp*, Supplementary Table S1) (35). We calculated GFP fluorescence density by dividing fluorescence (Supplementary Figure S4A) by FSC-A (Supplementary Figure S4B) measurements to account for cell size effects (Supplementary Results). Using an identical experimental setup as utilised to determine the loss of pH homeostasis, the *recA-gfp* strain was grown to the exponential phase before exposure to nalidixic acid in four different medias buffered to pHs 5.2, 6, 7 and 8 (Figure 2B). pH 6 promoted the most substantial expression of RecA-gfp while pH 8 showed the least expression. The FSC-A data also confirmed our previous findings and demonstrated the largest changes in cell size appear at the lowest pH after exposure to nalidixic acid (Supplementary Figure S4B).

These data illustrate that following DNA damage pH homeostasis is impaired and, consequently, affects RecA expression and downstream processes. These data also reinforce the fact that the external pH, via homeostasis loss, leads to altered morphological plasticity following DNA damage.

### Transcription of later induced SOS genes are repressed by an external pH of 5.2

Autoproteolytic degradation of the LexA transcription factor is a key step in the regulation and induction of the SOS response genes. To observe the induction of *lexA* an ectopic *lexA* promoter *gfpmut2* fusion was used (*plexA-gfp*, Supplementary Table S1) (36). The *plexA-gfp* system was investigated as outlined above for *recA-gfp*. The general pattern is similar to *recA-gfp* with a maximum expression around pH 6 with the following key differences: at pH 5 the *lexA* promoter activity is poorly induced and declines from180 min to 300 min Induction of the *lexA* promoter at pH 6-8 shows slightly sigmodial dynamics whereby induction accelerates to a maximum between 60-120 min whereas *recA* starts at maximum induction only slowing from 120 min (Figure 2C). Thus we show that *lexA* promoter expression is strongly influenced by the external pH with an optimal expression at pH 7.

The LexA protein is known to have asymmetric affinity to the heterogeneous SOS operator sequences and as such different genes are induced at varying LexA concentrations (37). As pH equilibration occurs gradually following DNA damage we hypothesised that the various SOS genes and their products could be differentially affected (38). PolV mediated mutagenesis is detrimental to the fitness of most bacteria and as a result the two SOS active components *umuD* and *umuC* of PolV are induced late in the SOS response (39). To monitor the induction of *umuDC* an ectopic *umuDC* promoter *gfpmut2* fusion was selected (*pumuDC-gfp*, Supplementary Table S1) (36). The *pumuDC-gfp* system was tested as outlined above for *recA-gfp*. The fluorescent signals of UmuDC-gfp start to increase after about 90 minutes of incubation, confirming *umuDC* is induced later than *recA*. Further, following nalidixic acid treatment *pumuDC* showed the highest and lowest induction in media at pH 6 and 5.2, respectively (Supplementary Figure S5A). Analysis of FSC-A in the *umuDC promoter* Gfpmut2 expressing cells at the indicated pH’s revealed the same filamentation patterns as *recA-gfp* expressing cells (Supplementary Figure S5B) confirming the previous findings (Figure 2A). After adjustment of the Gfpmut2 fluorescent intensity signals to cell size, *pumuDC-gfp* showed the strongest induction at pH 8 and practically no induction at 5.2 (Figure 2D), which is in contrast to *recA-gfp* (Figure 2B). To directly compare the pH dependency of early and late induced SOS genes, we plotted the 300 minutes after nalidixic acid exposure data points across the permissible pH range (Figure 2E-F). These data indicate early induced genes (*recA* and *lexA*) show maximum expression at pH 6 and 6.5 while late induced genes (*umuD* and *umuC*) showed maximum expression over a broader pH-range (6–8) (Figure 2D). Of the genes tested, only *recA* was found to be significantly induced at pH 5.2.

Finally, the identical experimental setup allowed temporal comparisons of pH equilibration against *recA-gfp, plexA-gfp* and *pumuDC-gfp* induction. Comparative fluorescence density measurements (Figure 3A) clearly confirmed the earlier induction of *recA-gfp* whose expression begins <30 minutes after exposure and up to 90 minutes before *umuDC-gfp* expression. The relative expression of *umuDC* is much faster and plateaus at 180 minutes, similar to *recA-gfp*. *plexA-gfp* shows an intermediate response, with a delayed start compared to *recA-gfp* but fast relative expression that plateaus after 180 min of exposure to the antibiotic. Comparison of intracellular pH and fluorescence density measurements also demonstrate induction of early-induced SOS genes such as *recA* and *lexA* occurs whilst the pH is at or still near physiological pH (Figure 3B-C). However, early expression for *recA* indicates that it is exposed to a large gradient of pH’s while later induced genes are expressed in an environment where the pH is already approaching that of the external environment (Figure 3D). Together, these results show that following DNA damage late induced SOS genes are more susceptible to a reduction or loss of induction at low external pHs whilst at pH 8 the opposite is observed.

**Figure 3.**
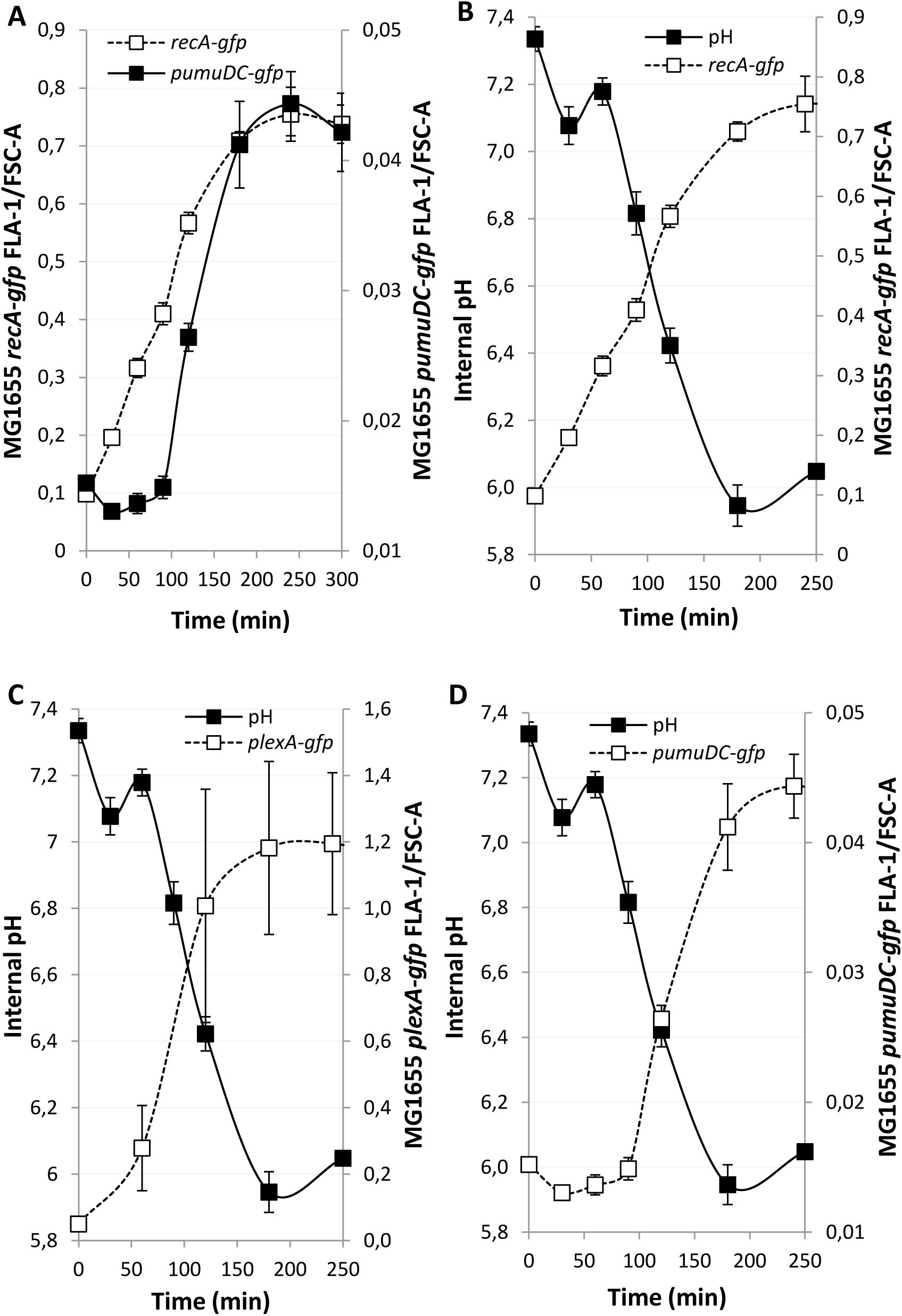
The contrasting induction kinetics of SOS genes leads to different pH environments for early and late genes. The media was buffered to pH 6 in all cases and DNA damage induced by nalidixic acid (100 µg/ml) at time point 0. All GFP signals were followed by flow cytometry at the indicated time points. (**A**) *recA* expression as determined by the fluorescence signal of *recA-gfp* is quickly induced following DNA damage as compared to *pumuDC-gfp*. (**B-D**) Intracellular pH and gene expression levels were inferred from the fluorescence signals of GFP. (**B**) The immediate induction of *recA-gfp* leads to significant induction occurring at homeostatic pH as compared to genes induced at later time points. (**C**) The kinetics of *plexA-gfp* induction leads to an intermediate situation where induction occurs mostly as the intracellular pH is falling. (**D**) The slower induction of *pumuDC-gfp* results in the induction occurring at non-homeostatic pHs.

### The biochemical activity of RecA is strongly dependent on pH

The strong correlation between RecA expression, the strength of SOS induction and their dependence on the external pH prompted us to investigate how pH affects known biochemical activities of RecA. First, we used an electrophoresis mobility shift assay to monitor the formation of RecA: DNA complexes. The binding of RecA to fluorescently labelled ssDNA is strongly pH dependent with almost 10-fold higher affinity observed at pH 5 compared to pH 8 (Figure 4A). This was accompanied also by a change in the mobility of the RecA-ssDNA complexes, indicating a possible conformational change of the filament (Supplementary Figure S6A). Second, a kinetic study using stopped-flow allowed us to further investigate the DNA binding properties of RecA at different pH’s by monitoring increasing fluorescence of a 5’Cy3-labelled dT79 oligonucleotide after binding with ATP pre-incubated RecA protein (Figure 4B-C and Supplementary Figure S6B) (20). Quantification of the binding parameters confirmed that decreasing pH led to higher affinity (Figure 4B) as well as a massive change in binding half-times (Figure 4C). Strikingly, the shift of pH from 5 to 8 resulted in a more than 3000 times increase in half-times. In other words, while at pH 5 it takes RecA less than one second to bind DNA, at pH 8 it needs several minutes. Finally, we used a D-loop formation assay to directly correlate the pH dependency of RecA-DNA binding with its ability to search for the homology within the donor dsDNA plasmid. Indeed, reactions at lower pH resulted in maximal activity, indicating that less RecA is required for the recombination reaction (Figure 4D and Supplementary Figure S6C). Based on altered SOS response we also elucidate previously reported RecA coprotease activity on LexA repressor (40). Our experiments show that RecA filaments stimulate LexA cleavage only at pH 6 in the presence of ATP (Figure 5A).

**Figure 4.**
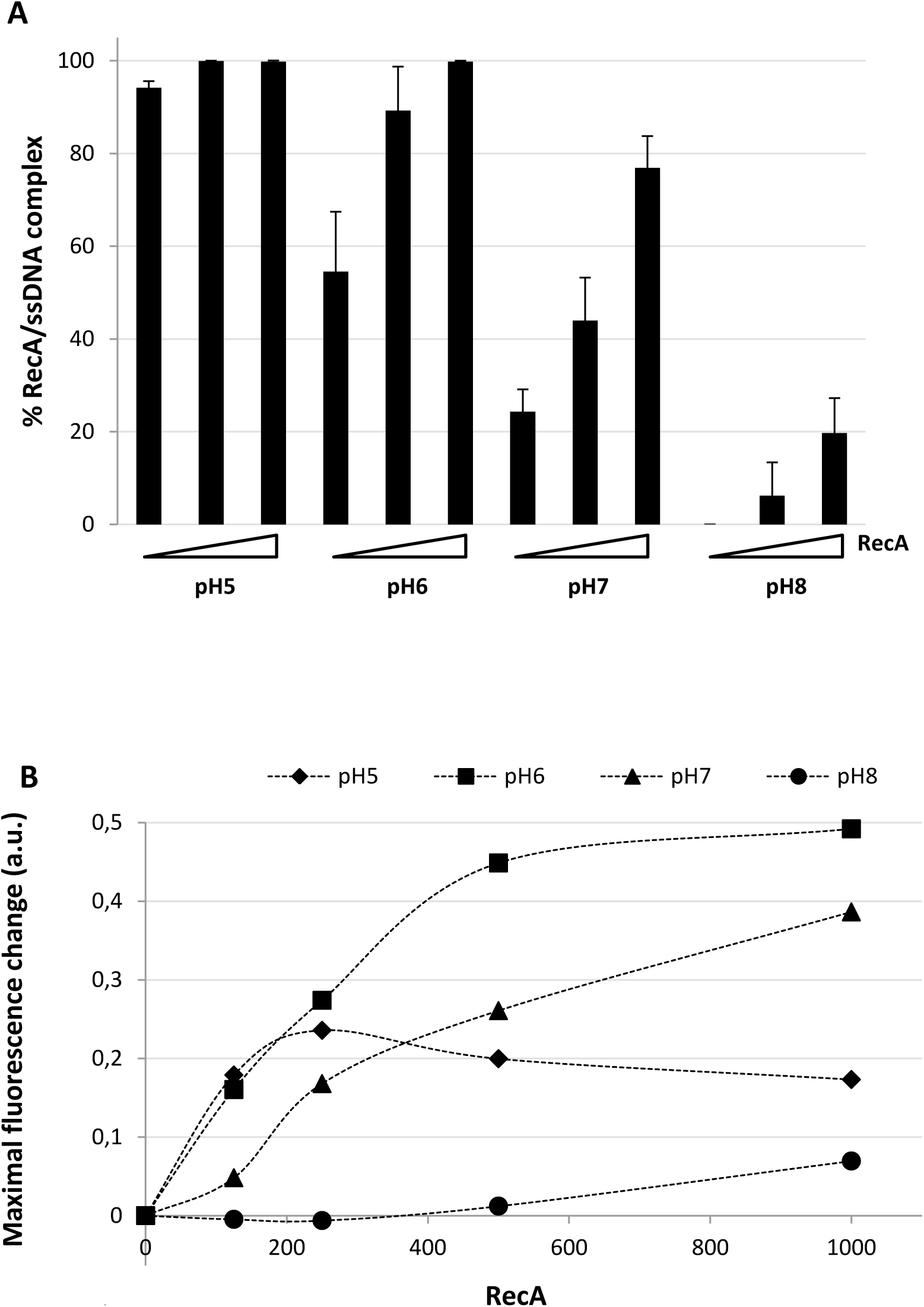
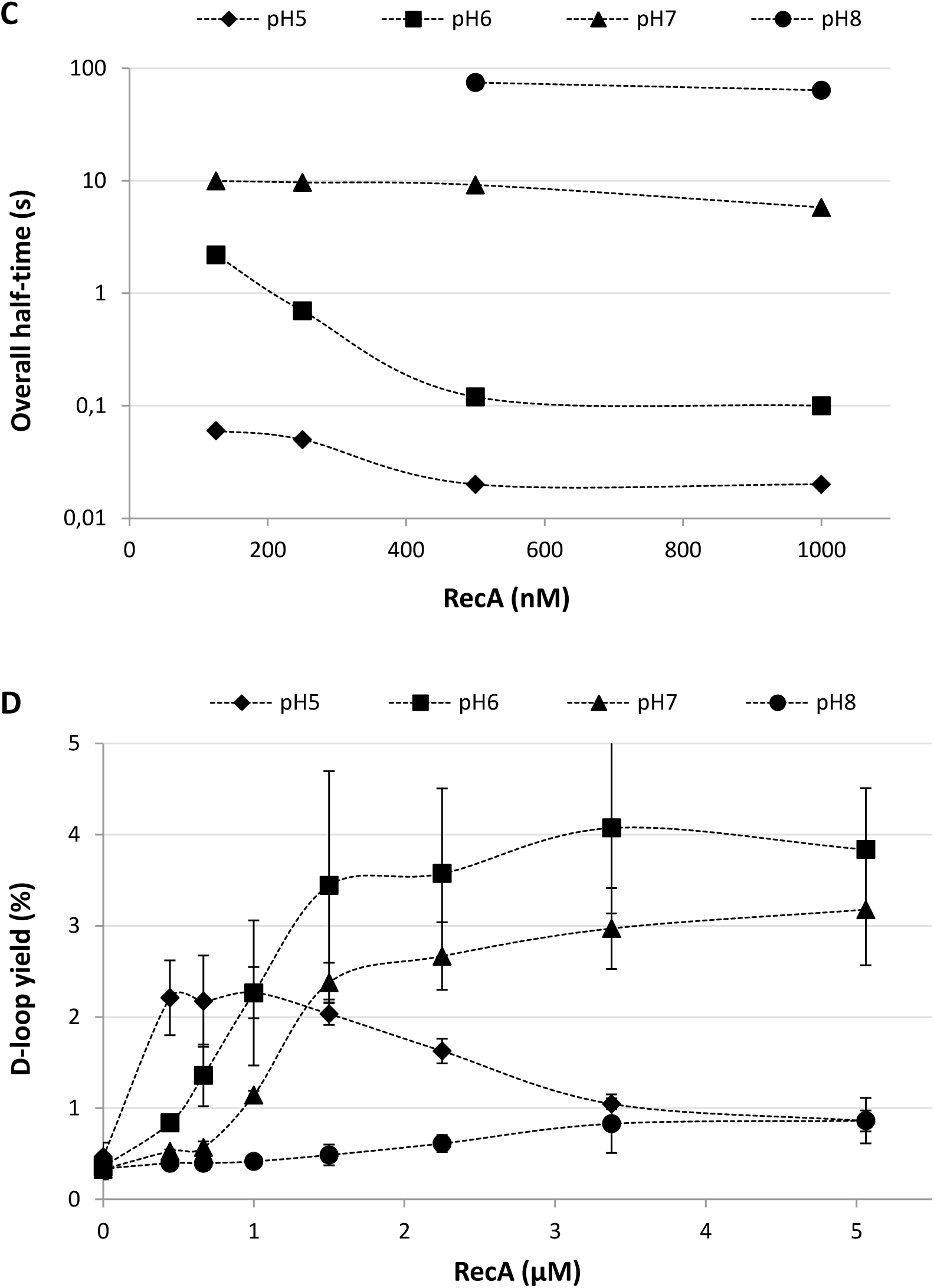
pH dramatically effects RecA biochemical activities. (**A**) The effect of pH on the ssDNA binding activity of RecA by EMSA. (mean ± s.d. error bars, (*n* = 3)). (**B**) pH dependency of RecA filament assembly by stopped-flow assay. Maximal change of fluorescence is depicted. (**C**) Overall half-times of ssDNA binding. (**D**) Efficiency of D-loop formation by RecA at various pH. (mean ± s.d. error bars, (*n* = 3)).

**Figure 5.**
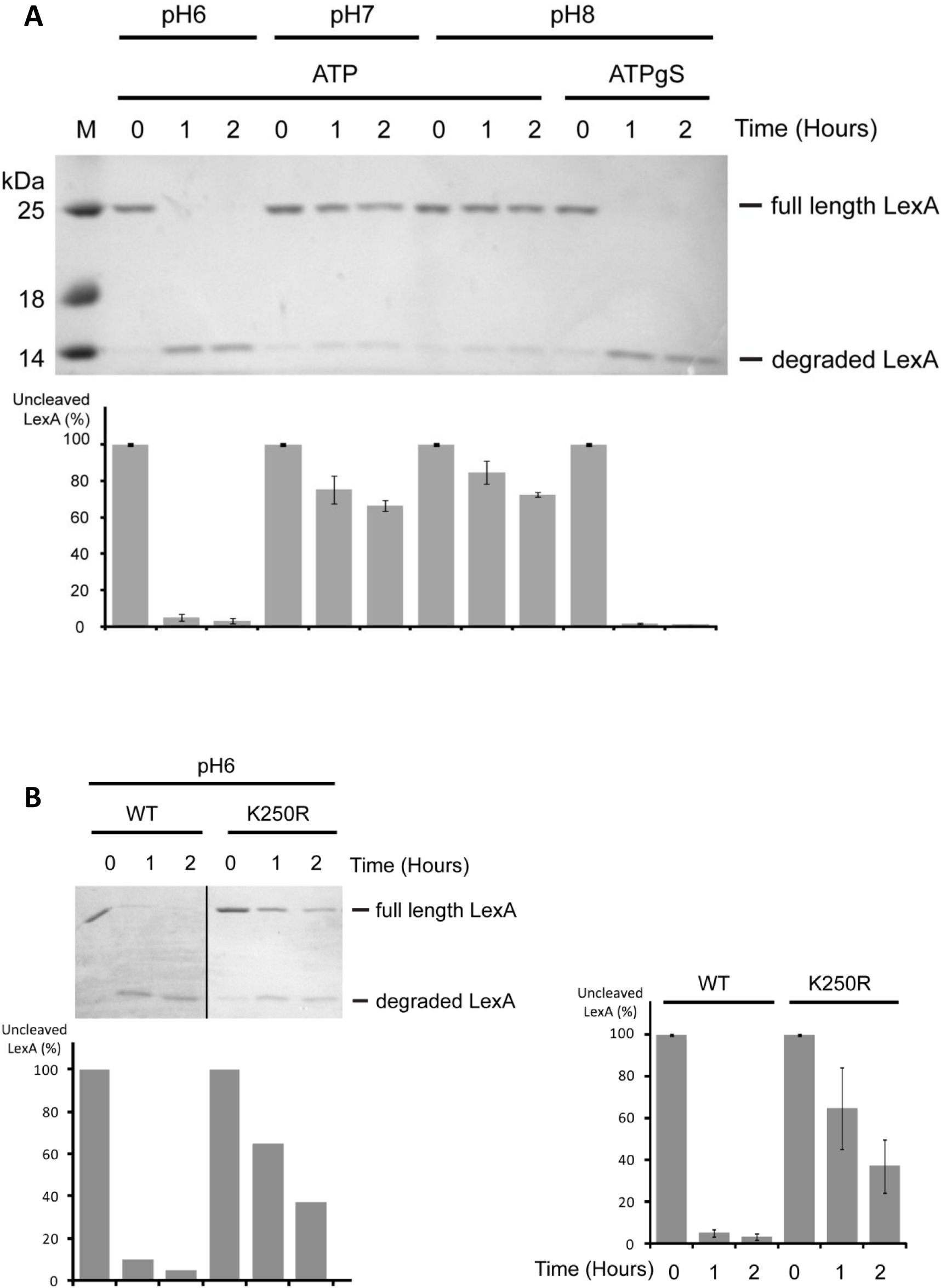
pH affects RecA co-protease activity on LexA cleavage. (**A**) The effect of pH on the RecA co-protease activity on LexA cleavage (mean ± s.d. error bars, (*n* = 3)) in the presence of ATP or ATPγS. (**B**) RecA K250R co-protease activity on LexA cleavage (mean ± s.d. error bars, (*n* = 3)) in the presence of ATP and pH 6.

To better understand the origins of transcriptional and RecA *in vitro* pH sensitivity we chose to examine RecA K250R a cationic mutant found at the subunit-subunit interface in RecA-ssDNA filaments that has shown pH dependent DNA three-strand exchange as well as a slow general recombination DNA repair phenotype (41). We hypothesised that this residue is responsible, at least partially, for the pH sensitivity we had observed. Initially we used RecA K250R to show a significantly impaired LexA autocatalytic activity (Figure 5B). As this mutant binds to ssDNA in a pH-dependent manner (Supplementary Figure S7), similar to the wt protein, it is plausible that impaired ATPase activity leads to a loss of coprotease activity. Taken together, these data show that pH strongly affects the biochemical activities of RecA and are remarkably similar to the *in vivo* SOS induction data.

### RecA K250 is important for SOS gene transcription at low pH

Subsequently, we wished to understand if *recA K250R* altered the observed pH depolarisation of the wild type strain. To do this the *recA K250R* strain (Supplementary Table S1) was transformed with the pH sensitive reporter to create *recA K250R GFPmut3\** and intracellular pH changes following antibiotic exposure were examined by flow cytometry (Figure 6A). We observed accelerated depolarisation kinetics compared to wild type *recA* where equilibration with the external media was achieved 1 hour earlier. This suggests that *recA K250R* was also having an effect on the SOS genes, and particularly those induced late in the response. Next we also monitored the SOS response of *recA K250R* using the GFP signal from the *pumuDC-GFP* plasmid (*recA K250R pumuDC-gfp*) following DNA damage by flow cytometry (Figure 6B). The transcriptional activity of *pumuDC* showed an increase in pH sensitivity as compared to wt *recA*, with the greatest difference between the two strains observed at pH 6-6.5 (Figure 6C), indicating that the K250 residue is responsible for increasing the transcriptional activity of *pumuDC* at lower pH.

**Figure 6.**
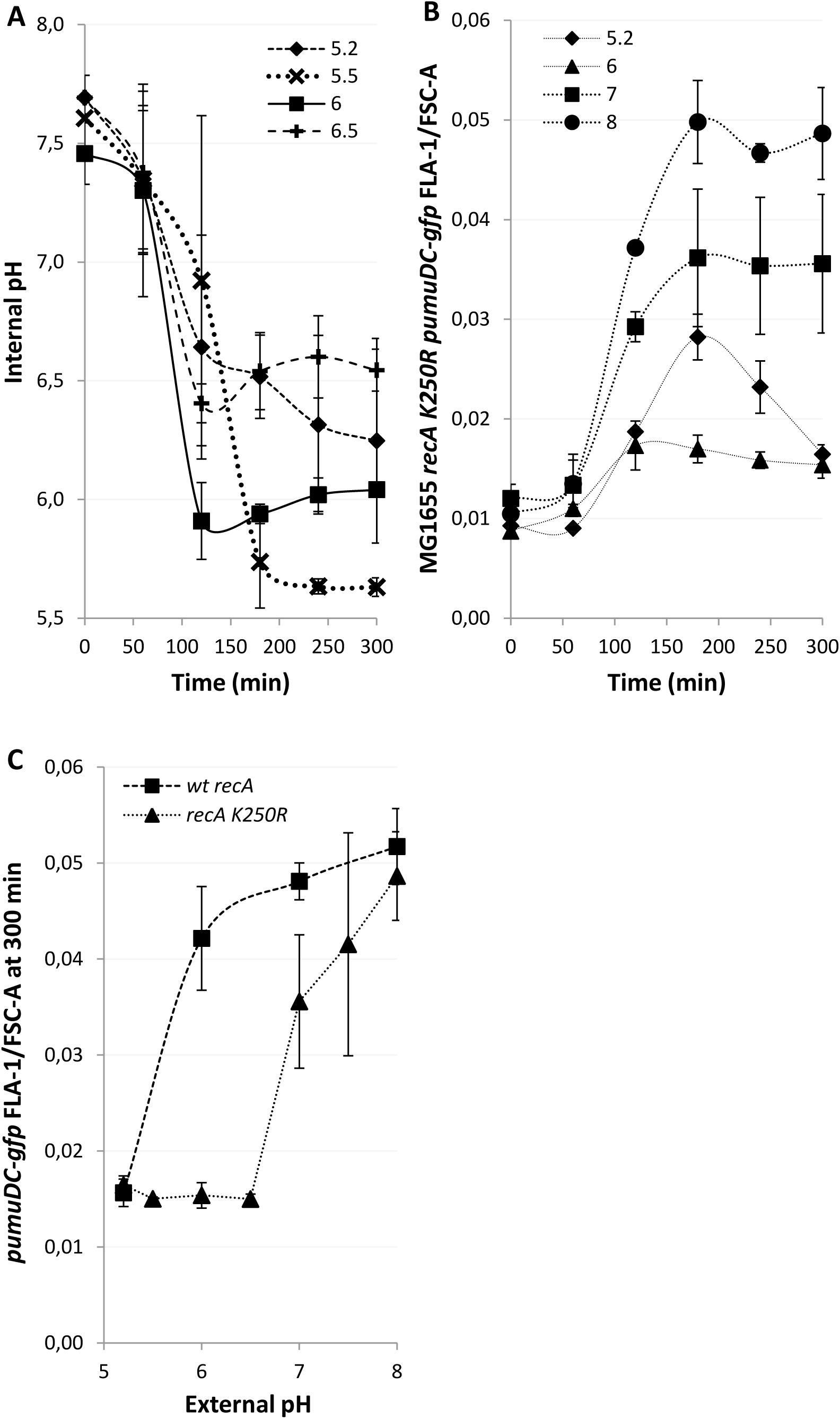
RecA K250 is responsible for increasing expression of late SOS genes at low pH. All GFP signals were recorded using a flow cytometer. (**A-B**) DNA damage was induced at time point 0 with nalidixic acid (100 µg/ml) and the GFP signals recorded at the time points indicated. (**A**) pH equilibration with the external media over the viable *E. coli* range is faster with RecA K250R than wt RecA. (**B**) The late induced operon *umuDC* shows reduced expression at lower external pH’s in a *recA* K250R background. (**C**) The differences in expression are greatest at pH 6-6.5 where virtually no expression is seen in the *recA* K250R background following 5 hours of exposure to Nalidixic acid. For each data point the mean ± s.e.m. error bars is shown, (*n* = 3 independent replicates))

### Mutagenesis is strongly influenced by pH

DNA damage-induced loss of pH homeostasis and corresponding changes in RecA activities prompted us to hypothesise that genome stability may be affected over the range of studied pH. The rate of mutagenesis was initially monitored using a lactose reversion assay where the media was buffered across the permissible replicative range in the *E. coli* FC40 strain (Supplementary Table S1) (42). Since the lactose reversion assay detects frameshift mutations, base substitutions, and gene amplifications/ genome rearrangements, which in turn are dependent on a variety of SOS genes including *recA, recBC, dinB, polB, and umuDC* (43), this allowed us to also monitor the cumulative effect of the individual pathways in mutagenesis. The data illustrates the strong influence external pH has over the rate of mutagenesis (Figure 7A). The frequency of lactose reversions is close to zero at the extremes, pH 5 and 8, but reaches a maximum between pH 6.5 and 7 (Figure 7B). Colonies formed at pH 7.5 arise due to lac-amplification rather than mutations as they occurred only gradually. In contrast, most of the colonies at pH 6.5 and 7 form relatively quickly over the first few days, suggesting point mutations are the predominant mechanism (44). These data demonstrate how the external environment could affect the rate of mutagenesis in niches of different pH.

**Figure 7.**
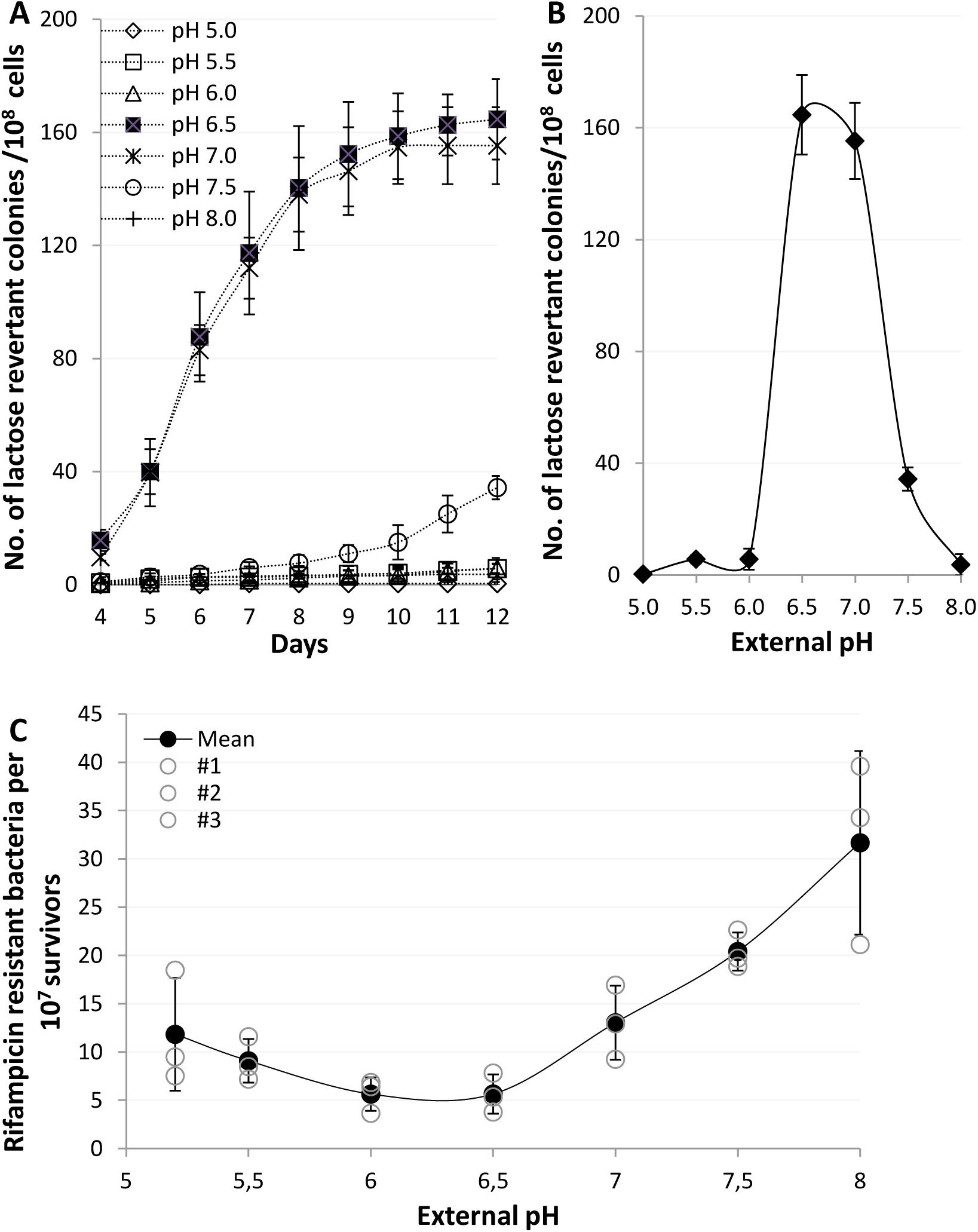
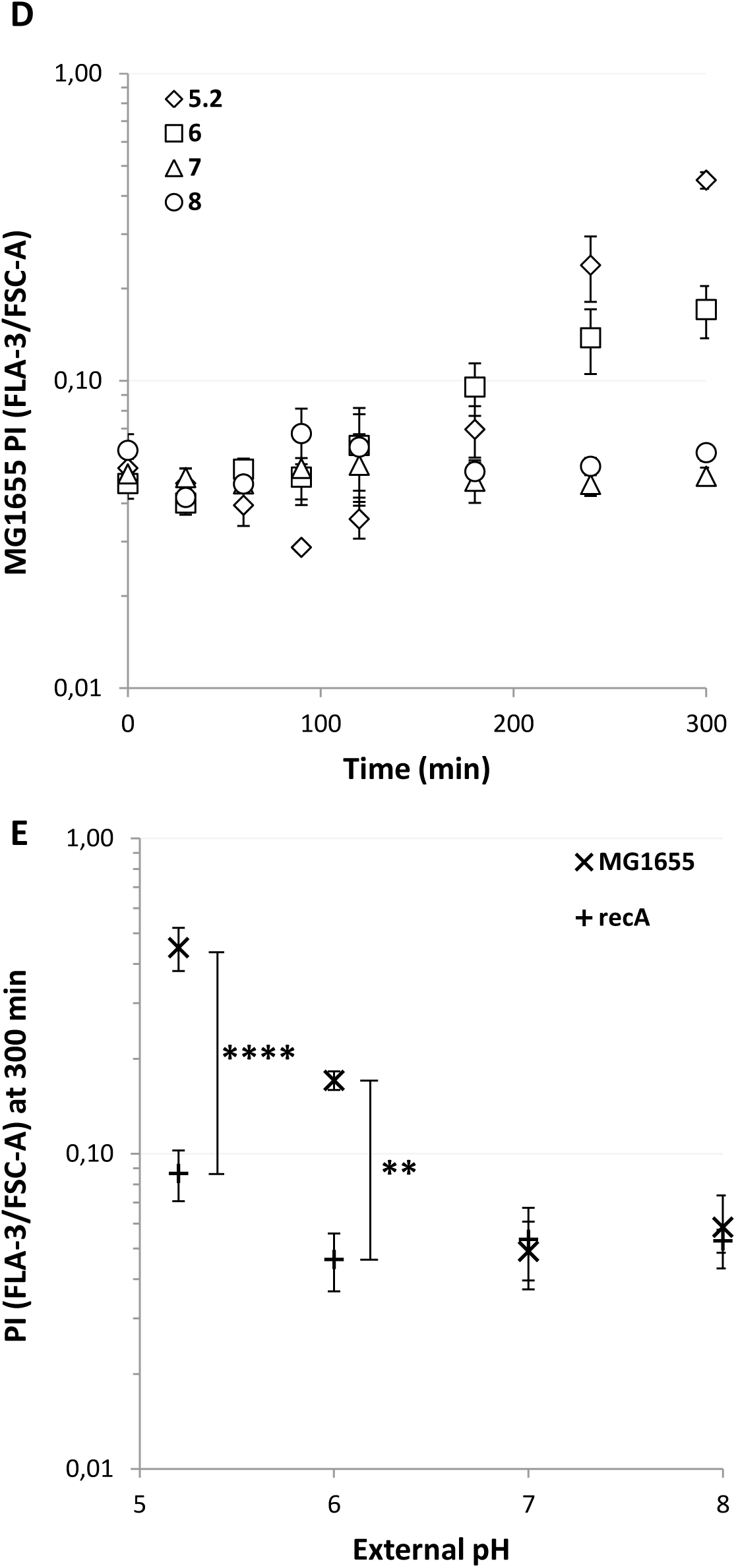
Extracellular pH strongly influences mechanisms of genetic resistance development and membrane integrity. (**A-C**) For each data point the mean ± s.e.m. error bars is shown, (*n* = 3 independent replicates). (**A**) A lactose reversion assay using *E. coli* strain FC40, which cannot utilize lactose, when grown on minimal media plates with lactose as the sole carbon source. Genetically resistant, reverted, populations appear over several days and the rate is indicative of the rate of resistance development. (**B**) A pH dependence profile after 12 days showing the total number of lactose reverted colonies. (**C**) pH dependence of a forward UV induced rifampicin resistance assay. Mean, filled circles ± standard deviation (n=3 (independent replicates), empty circles). (**D-E**) DNA damage was induced by nalidixic acid (100 µg/ml). For each data point the mean ± s.e.m. error bars is shown, (*n* = 4 independent replicates) (**D**) The more acidic the environment the higher the level of membrane instability and permeability. PI density staining following DNA damage induction with nalidixic acid at time point 0 at the pH’s and time points shown. (**E**) PI density staining after 5 hours of exposure to nalidixic acid shows that the membrane instability at acidic pH’s is *recA* dependent. ** p ≤0.01, **** p ≤0.0001 (t-test, two tail, unpaired equal variance).

Subsequently, we examined the pH dependency of a forward mutation assay as opposed to the lactose reverse mutation assay. We used UV to directly induce DNA damage and increase mutagenesis (Figure 7C), since all antibiotic minimum inhibitory concentration assays tested showed pH dependency (Figures S8A-H). In this case mutagenesis showed pH dependency with an observed minimum at pH 6 and a maximum at pH 8. We concluded that mutagenesis is significantly influenced by pH but that the specific stimuli the bacteria are exposed to alter the response. In our examples, the lactose reversion assay, which is dependent on both entering into the stationary phase and the DNA damage response results in the highest mutational frequency from pH 6.5-7 whilst when only the SOS response is required the lowest frequencies were found at pH 6.

To explain the differences in mutagenesis over the viable pH range for *E. coli* we hypothesised that bacterial viability was pH dependent. We used propidium iodide (PI) staining to monitor bacterial viability whereby PI enters the bacteria only upon loss of membrane integrity. The results showed that the bacteria lose membrane integrity at acidic pH after nalidixic acid exposure, whereas at pH 7-8 the membranes of the bacteria remain impermeable to PI (Figure 7D). An identical experiment using a *recA* mutant showed the loss of membrane permeability at low pH was significantly dependent on *recA* (Figure 7E). Strikingly, at pH 8, where we recorded the highest number of rifampicin resistant mutants there was both the lowest PI staining and highest transcription of *pumuDC-gfp.* Additionally we investigated viability using a classical colony forming survival assay (Supplementary Figure. S9), however, we observed large variations at and after two hours that prevented any interpretation of the significance of external pH.

Together our results clearly demonstrate that the environmental pH where bacteria and antibiotics meet plays a significant role in antibiotic resistance development. This appears to occur due to proton equilibration across the plasma membrane so that the intracellular pH becomes that of the environment. This pH change subsequently affects many downstream processes of the SOS DNA damage response system ultimately leading to pH-dependent mutagenesis.

## Discussion

In this work we aimed to understand the physiological consequences of environmental pH on the SOS response. We initially discovered, in response to the DNA-damaging antibiotic nalidixic acid, a failure of pH homeostasis in the bacterial population. This failure occurred in a controlled fashion over several hours as the cytoplasmic pH equilibrated with the extracellular pH environment. Subsequently, we showed the pH of the external media affects the strength of induction of the SOS genes *recA*, *lexA* and *umuDC*, and the rate of mutagenesis resulting in antibiotic resistant mutants. We also examined how pH affects the biochemical function of the RecA protein *in vitro* and show that at pH 8 binding of ssDNA was severely compromised, a finding which correlates well with the *in vivo* weak induction of *recA.* Additionally, we found that RecA K250, a pH sensitive residue in RecA, was partially responsible for RecA inducing *pumuDC-gfp* expression and essential up to a pH of 6.5. Taken together our results indicate that the pH of the surrounding environment can influence and enhance mutagenesis and thus accelerate antibiotic resistance development (Figure 8). We also suggest that pH depolarisation across the inner membrane could be a mechanism whereby bacteria directly sense the environmental pH following a lethal challenge to their DNA integrity. By this mechanism the environmental pH would automatically be sensed by all intracellular components including all translated proteins.

**Figure 8.**
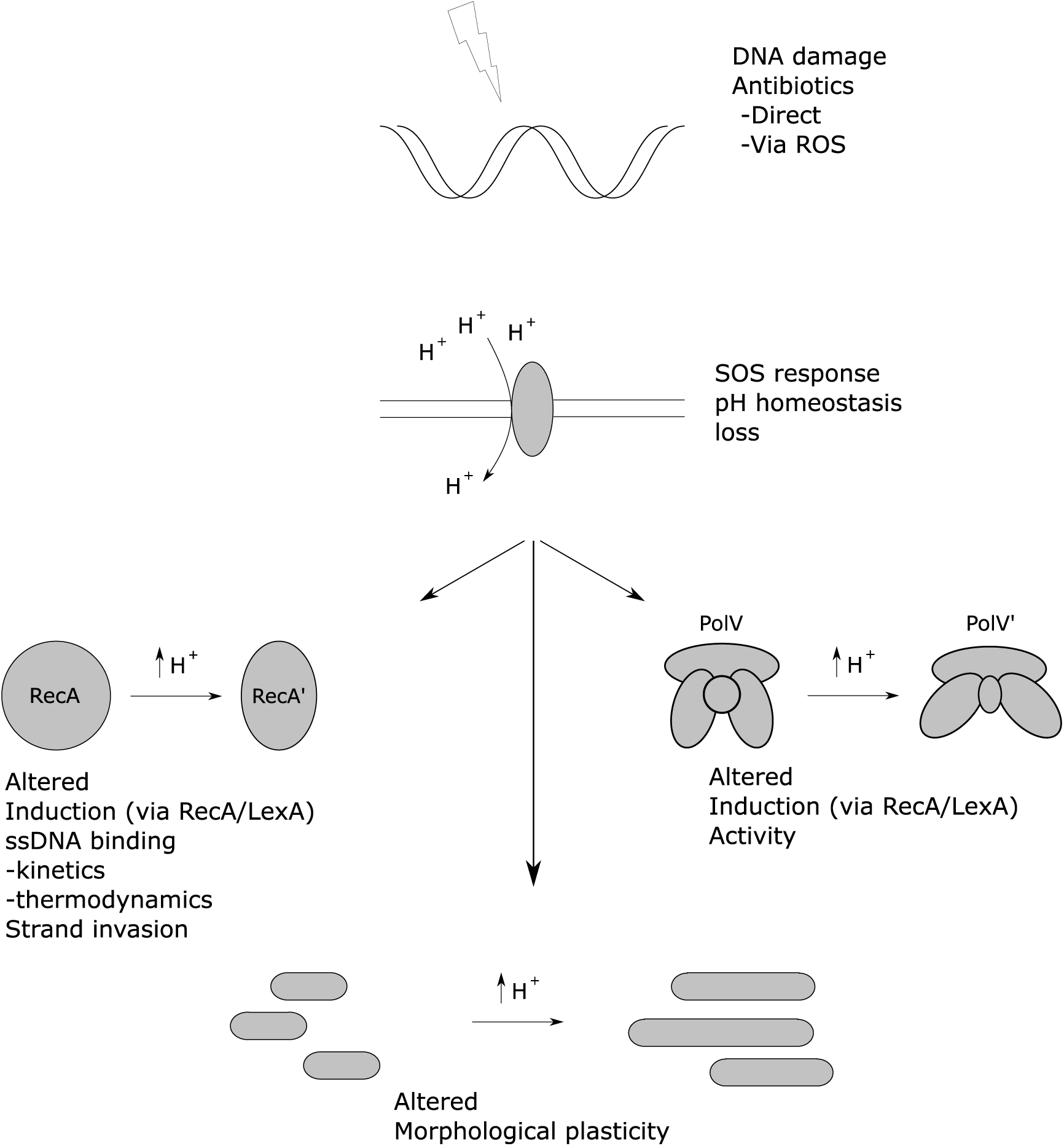
Proposed working model of the flow of pH dependent events following DNA damage. DNA damage, for example from exposure to antibiotics, results in the stimulation of the SOS response. The SOS response results in a gradual loss of pH homeostasis, in turn the proton depolarisation results in an altered biochemical environment touching all biological processes. Here we specifically show *recA* induction, RecA’s kinetics and thermodynamics with regards to ssDNA and strand invasion are affected. Additionally morphological plasticity and other downstream processes such as the activity of TLS polymerases are affected.

### Environmental pH influences antibiotic resistance development

Specifically, our findings show the following pH-dependent responses to nalidixic acid treatment: At an external pH of 5.2 genetic instability is not detectable. This could be explained by the lack of *pumuDC* transcription and significant loss of membrane integrity. The primary survival response at this pH appears to be filamentation. As the external pH increases, membrane integrity improves so that it is roughly the same as before the antibiotic was applied at pH’s 7 and 8, at the same time the transcriptional activity of the genes studied increases, increasing the probability that translesion polymerases will be active. This could explain the increased genome instability seen at the higher pH’s. These increases in genome instability at pH’s above 6.5 also imply that the development of resistance to antibiotics will be increased under these circumstances.

### The activities of RecA are sensitive to pH

Early transcriptional activity leads RecA to be exposed to all possible pH’s from homeostatic to environmental pH. Previous *in vitro* studies have shown pH 6.2 is optimal for stable RecA filamentation on ssDNA (45). Experiments comparing the binding of SSB protein and RecA on ssDNA at pH’s 7.5 and 6.2 showed the lower pH greatly favoured RecA over SSB (46). DNA independent hydrolysis of ATP was significant at pH 6 whilst undetectable at pH 7 (47). More recent work has demonstrated both RecA nucleation and subsequent filament extension on SSB-ssDNA complexes are strongly pH dependent, where pH 6.5, the lowest examined pH, gave the highest nucleation and filamentation rates (48). We also found pH dependencies regarding RecA’s interaction with ssDNA. Affinity was strongest and rate of binding fastest at pH 5 and became respectively weaker and slower as pH increased. Ultimately we discovered almost no D-loop formation at pH 8 with the implication that recombination is unlikely to occur *in vivo* once the intracellular pH reaches 8. Optimal D-loop formation was seen at pH’s 6-7 indicating that recombination could potentially be an important resistance development mechanism at these pH’s. The origins of the pH sensitivity have been postulated to involve an electrostatic interaction between the negatively charged core domain of one RecA with the positively charged N-terminal domain of the next (49). Accordingly, we examined the effect of the K250R RecA residue predicted to be involved in inter-protein interactions. We found this mutant to be highly influenced by pH, this was most clearly evident at pH 6 when compared to the wild-type, here there was a significant reduction in LexA degradation and no expression of *umuDC*. We conclude, therefore, that K250 and monomer-monomer interactions are important at low pH and influential up to pH 7.5.

### A controlled loss of pH homeostasis

The kinetic profile of proton depolarisation after DNA damage indicates how the mechanism may proceed. The observed process contrasts with a physical breach of the membrane which results in a near instantaneous pH equilibration. Indeed, even a rapid shift in external pH without exposure to DNA damaging agents, for example, a shift from 7.5 to 5.5 can result in an intracellular pH shift immediately to 6, with a biphasic recovery beginning after 10 seconds and most of the physiological pH recouped after 30 seconds, the remaining being recovered over the next 4 minutes (24). Based on our data we concluded, therefore, that the observed mechanism did not involve any rapid disruption of the inner membrane, temporary or permanent, but a gradual and controlled loss of pH homeostasis.

The bacterium expends considerable resources in maintaining pH homeostasis and optimal internal environment for the metabolism of the cell (50). However, our data point to situations where the loss of homeostasis and thereby direct environmental sensing of pH may be an advantage. As such the pHs used for the study of bacterial proteins outside of the homeostatic range of 7.5-7.7 may be biologically relevant in such cases. In an optimal environment the maintenance of ion gradients, to generate a pmf, is important for energy production. However, under DNA damaging conditions, it could be an advantage for the cell to enter a state of persistence, a dormant condition where energetic processes are reduced to a minimum. Recently, dependency between intracellular ATP concentrations and persister cell formation has been established (51). Following DNA damage we show that, in addition to the loss of ΔΨ, the cell also suffers a loss of ΔpH thus greatly reducing the capacity of the bacterium to generate ATP.

### Dynamic bacterial morphology following filamentation

We found the reduction in bacterial filamentation after 1 hour of exposure to Nalidixic acid surprising as we assumed that filamentation would continue until either the DNA damage stress is removed or the bacterium succumbs to the stress. Yet, recent work suggests cell division could be a response to severe DNA damage-induced stress. In order for cell division to occur replication must be completed. When irreparable damage occurs on one DNA strand this dooms the bacterium that receives this permanently damaged chromosome but allows the other bacterium with an undamaged chromosome to continue proliferation (52, 53). Double strand breaks as produced by inhibition of topoisomerase ligation activities as in the case for nalidixic acid could lead to a serious impact on bacterial viability for both chromosomes. Yet it was interesting to note that reducing cell size following filamentation correlated with the start of the slow phase of bacterial killing attributed to the presence of persister cells in response to an overwhelming challenge to DNA integrity. We see this time point as a change of survival tactics to better suit a chronic challenge to DNA integrity.

### A low external pH environment leads to enhanced filamentation with implications for phagocytosis

Varying intracellular pH should lead to altered biochemical activities of the cellular proteins. Filamentation, or morphological plasticity, is a physiological change seen in bacteria in response to DNA damage. The mechanism involves many components but initiation is caused by the SOS protein SulA. Specifically SulA binds FtsZ and thereby inhibits polymerisation of FtsZ which would otherwise form a ring at mid-cell and thus inhibits septation (54). Other constituents involved in filamentation are proteins participating in cell membrane and cell wall synthesis. From our results, it is clear the sum of the pH effects on these components together leads to longer filaments at lower pH, specifically 5.2 and 6 whilst at pH 7 and 8 the filaments are shorter. It is interesting to note in uropathogenic *E. coli* SulA is essential for pathogenesis in immunocompetent hosts (55, 56). It has been postulated an influx of neutrophils following bacterial emergence from epithelial cells results in the phagocytosis of bacillary forms of uropathogenic *E. coli* but not filamented forms. Filamented *E. coli* appear to be resistant to phagocytosis and exposure to an acidic environment, such as in macrophages leads to a stronger filamentation phenotype so if the bacterium survives it is unlikely to be engulfed again.

In conclusion we find that the pH of the environment will significantly affect the mechanisms of antibiotic resistance and the rate at which they occur in response to antibiotic treatment.

## Acknowledgements

Acknowledgements and Funding

We are indebted to D.B. Weibel, M. Rajendram, J. L. Slonczewski, S. Rosenberg, M.M. Cox, U. Alon & J. Stavans for providing strains and plasmids. We also thank M.M. Cox for the RecA K250R protein. This work was supported by Czech Science Foundation (GACR 13-26629S and 207/12/2323); project no. LQ1605 from the National Program of Sustainability II (MEYS CR), FNUSA-ICRC no. CZ.1.05/1.1.00/02.0123 (OP VaVpI), Trond Mohn Foundation (Norway) TMS2019TMT05 and South-Eastern Norway Regional Health Authority, Innovation Fund 30279.

