## Supplementary files for "Antibiotic-induced DNA damage results in a controlled loss of pH homeostasis and genome instability"

### Supplementary Results

#### DNA damage leads to the collapse of the electrical potential

Electrical depolarisation is a hallmark of the SOS response (1,2). Our initial step was to confirm previously observed changes in electrical polarity of *E. coli* following DNA damage. To induce DNA damage and the SOS response the bacteria were exposed to the quinolone antibiotic nalidixic acid (3). The depolarisation kinetics were similar to those previously reported, an initial delay of 60 minutes is seen before a steady depolarisation over the next 240 minutes (Supplementary Figure S1)(1,2). The two key upstream regulators in the activation of the SOS response are the single-stranded DNA (ssDNA) binding protein RecA and LexA the RecA-ssDNA nucleoprotein sensitive repressor of the SOS genes. To confirm the SOS response dependency on electrical depolarisation an identical experiment was carried out with a *recA* mutant (Supplementary Figure S1).

#### GFP fluorescence is pH dependent

In order to measure the intracellular pH of bacteria, we used the pH sensitive properties of a fast folding derivative of GFP, GFPmut3* (4). Although convenient, observations from GFP based reporter systems should take account of filamentation that occurs during SOS. The fluorescence signal of GFP and other fluorophores have been demonstrated to correlate linearly with cell size and do not necessarily show an increased response due to their underlying inducer (5,6). Cell size can be defined by forward scatter (FSC) during flow cytometry, however, side scatter (SSC) has also been shown to be an equally valid measure (5,6). Subsequently, observations of fluorophore signal intensities when cell filamentation is observed have used FSC to correct for changes in fluorescence due to cell morphology (7). The pH sensitivity of a GFPmut3* containing system was assessed using the membrane permeable weak acid sodium benzoate that rapidly equilibrates the pH across the inner membrane within the permissive range of pHs for neutrophilic *E. coli* 5.5-8 (Supplementary Figure S2). To improve accuracy and to take account of changes in morphology and other possible factors affecting fluorescence a calibration curve was produced for each data point (Supplementary Figure S3). In this example, the calibration curves are biphasic with an increase in fluorescence signal for the first 90 minutes followed by a reduction for the remainder of the time. Together these experiments illustrate how the use of calibration curves for each data point can aid in the production of accurate data.

**Table S1. Strains and plasmids used in this study**

| Strain | Genotype / plasmid | Parent strain | Source |
|---|---|---|---|
| MG1655 | *F-, λ-, rph-1* | - | |
| **GFPmut3*** | */ pAra TorA-GFPmut3** | MG1655 | Electroporation of plasmid from (4) |
| *ΔrecA* | *ΔrecA* | MG1655 | P1 transduction from JW2669-1 (8) and pCP20 removal of Kan$^R$ |
| ***ΔrecA* GFPmut3*** | */ pAra TorA-GFPmut3** | MG1655 *ΔrecA* | Electroporation of plasmid from (4) |
| *recA-gfp* | *ygaD1::kan recAo1403 recA4136::gfp-901* (*gfp901* refers to *mut-2*) | MG1655 | (9), the original genotype SS3041 was from here (10) |
| ***p**lexA-gfp* | */ plexApromoter-gfpmut-2* | MG1655 | Electroporation of plasmid from (11) |
| ***p**umuDC-gfp* | */ pumuDCpromoter-gfpmut-2* | MG1655 | Electroporation of plasmid from (11) |
| *recA K250R* | *recA K250R* | MG1655 | P1 transduction from EAW540† (a generous gift from M. M. Cox) and pCP20 removal of Kan$^R$ |
| ***recA K250R* GFPmut3*** | */ pAra TorA-GFPmut3** | MG1655 *recA K250R* | Electroporation of plasmid from (4) |
| ***recA K250R pumuDC-gfp*** | */ pumuDCpromoter-gfpmut-2* | MG1655 *recA K250R* | Electroporation of plasmid from (11) |
| SMR4125 (FC29) | *Δ(lac-proAB)*$_{XIII}$ *ara thi* Rif$^S$ *[F' proAB$^+$ Δ(lacI-lacZ)]* | P90C | (12) |
| SMR4562 (FC40) | *Δ(lac-proAB)*$_{XIII}$ *ara thi* Rif$^R$ *[F' proAB$^+$ lacI33ΩlacZ]* | P90C | (13) |

†EAW540 contained an mutant FRT-Kan$^R$-wt FRT just after the stop codon of *recX* for transduction and *recA K250R*.

**Figure S1**

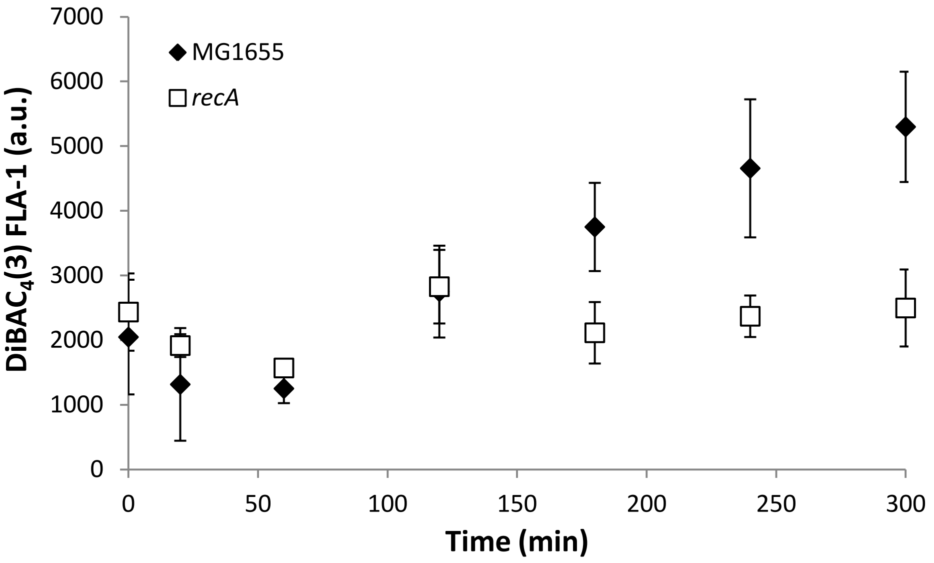

MG1655
recA
DiBAC$_4$(3) FLA-1 (a.u.)
7000
6000
5000
4000
3000
2000
1000
0
0
50
100
150
200
250
300
Time (min)

**Figure S1. DNA damage results in *recA* dependent depolarisation.**

The polarity of the bacteria was followed using the negatively charge dye DiBAC$_4$(3) that increases in fluorescence upon entry into the bacterium. DNA damage was induced in an exponentially expanding population of *E. coli* (MG1655 or MG1655 *recA*) by addition of nalidixic acid (100 µg/ml) at 0 minutes and the fluorescence intensity of DiBAC$_4$(3) followed using flow cytometry at the time points indicated. Plotted points are mean ± s.d. (*n* = 3).

**Figure S2**

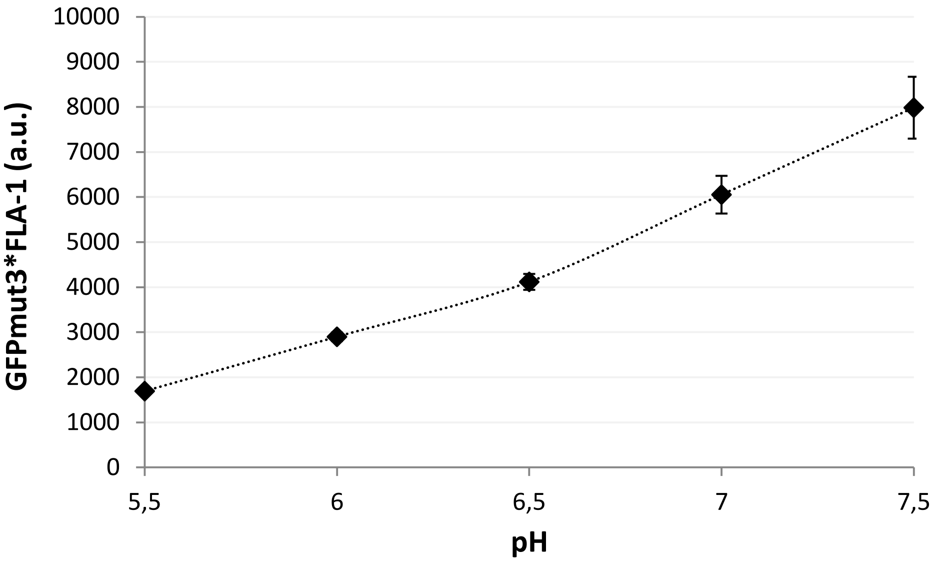

10000
9000
8000
7000
6000
5000
4000
3000
2000
1000
0
GFPmut3*FLA-1 (a.u.)
5,5
6
6,5
7
7,5
pH

**Figure S2. Fluorescence intensity of intracellular GFPmut3* varies according to pH.**

Calibration of GFPmut3* using buffered media at the pH's indicated and equilibrated with sodium benzoate in an exponentially expanding suspension of *E. coli*. No DNA damage was induced, fluorescent intensity values were measured by flow cytometry. Plotted points are mean ± s.d. ($n$ = 4).

**Figure S3.**

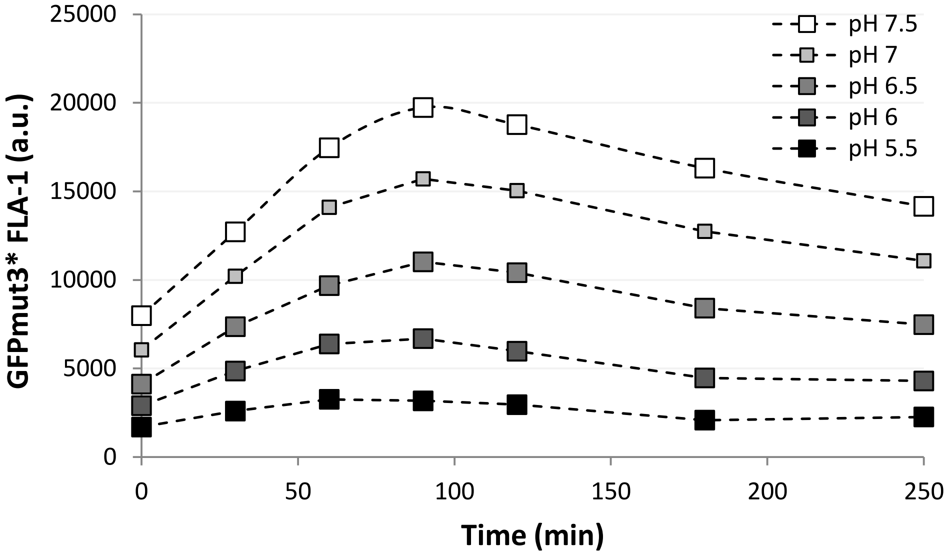

25000
20000
15000
10000
5000
0
0
50
100
150
200
250
GFPmut3* FLA-1 (a.u.)
Time (min)
pH 7.5
pH 7
pH 6.5
pH 6
pH 5.5

**Figure S3. Temporal fluorescence intensity measurement variation following DNA damage results in fluctuations in the calibration curves,**

Calibration of GFPmut3* using buffered media at the pH's indicated following equilibration with sodium benzoate. DNA damage was induced in an exponentially expanding population by nalidixic acid addition (100 µg/ml) at t=0, fluorescence intensity values were measured by flow cytometry. Plotted points are mean ± s.d. (*n* = 4).

**Figure S4.**

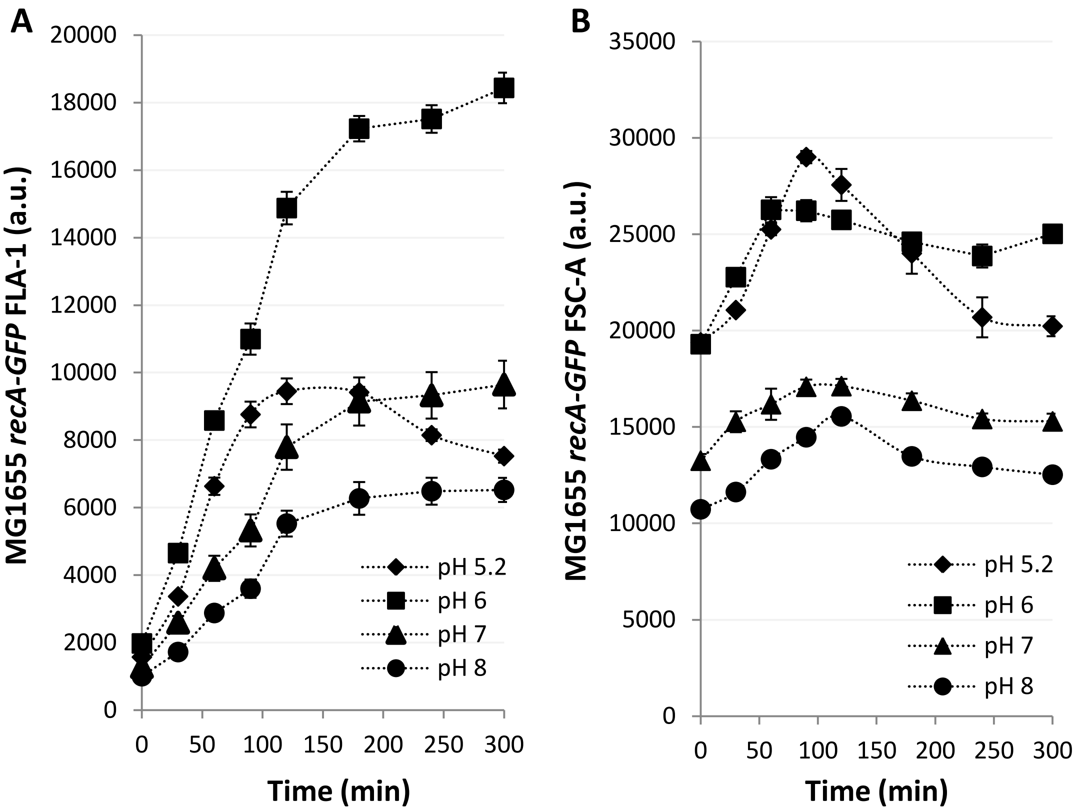
A
20000
18000
16000
14000
12000
10000
8000
6000
4000
2000
0
0 50 100 150 200 250 300
Time (min)
MG1655 *recA-GFP* FLA-1 (a.u.)
pH 5.2
pH 6
pH 7
pH 8
B
35000
30000
25000
20000
15000
10000
5000
0
0 50 100 150 200 250 300
Time (min)
MG1655 *recA-GFP* FSC-A (a.u.)
pH 5.2
pH 6
pH 7
pH 8

**Figure S4.** ***recA*** **transcription and cell morphology are dependent on the external pH following DNA damage**

(**A**) The fluorescence signal of *recA-gfpmut2* was followed using flow cytometry.

(**B**) Morphological remodelling was followed by flow cytometry using FSC-A as a proxy for bacterial size.

(**A & B**) The liquid media was buffered to pH 5.2, 6, 7 and 8. DNA damage (nalidixic acid 100 µg/ml) was induced at t=0 min and error bars are mean ± s.e.m., $n$ =4 (independent replicates).

**Figure S5.**

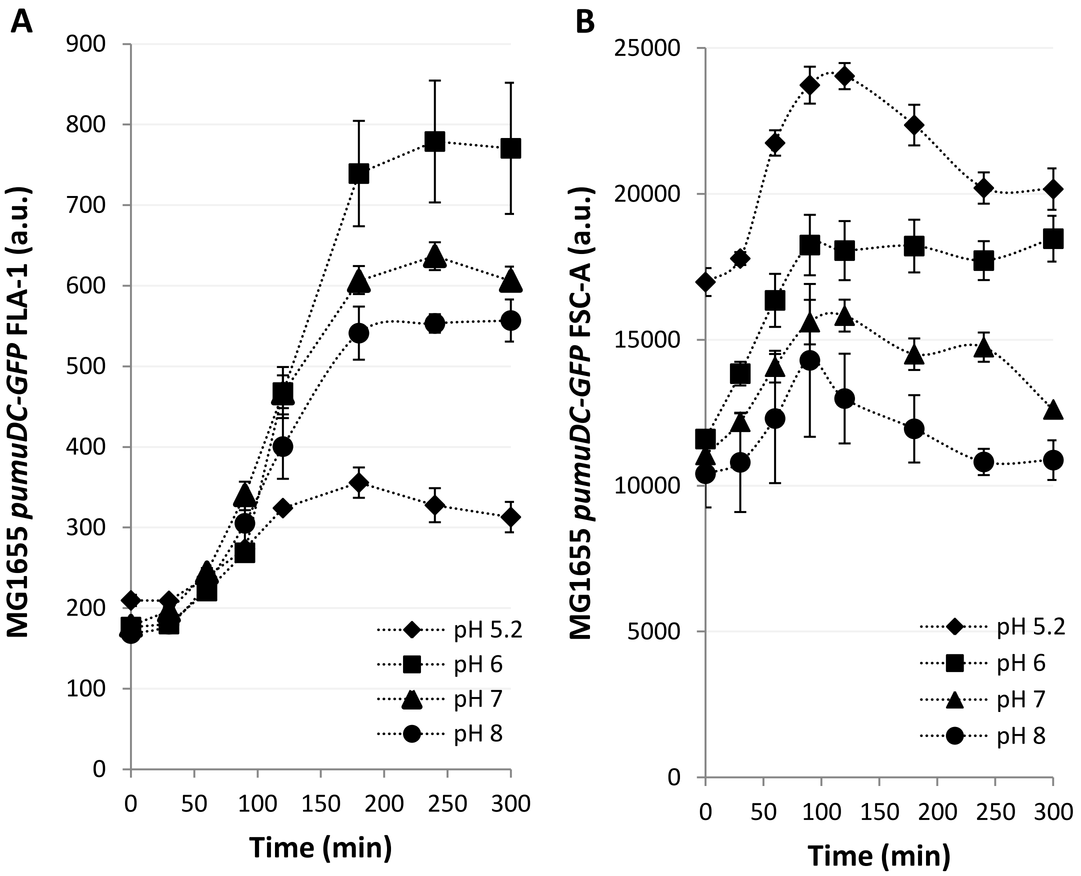
A
MG1655 *pumuDC-GFP* FLA-1 (a.u.)
Time (min)
pH 5.2
pH 6
pH 7
pH 8
B
MG1655 *pumuDC-GFP* FSC-A (a.u.)
Time (min)
pH 5.2
pH 6
pH 7
pH 8

**Figure S5.** ***umuDC*** **transcription and cell morphology are dependent on the external pH following DNA damage**

(**A**) The fluorescence signal of *pumuDC-gfpmut2* was followed using flow cytometry.

(**B**) Morphological remodelling was followed by flow cytometry using FSC-A as a proxy for bacterial size.

(**A & B**) The liquid media was buffered to pH 5.2, 6, 7 and 8. DNA damage (nalidixic acid 100 µg/ml) was induced at t=0 min and error bars are mean ± s.e.m., $n$ =4 (independent replicates).

**Figure S6.**

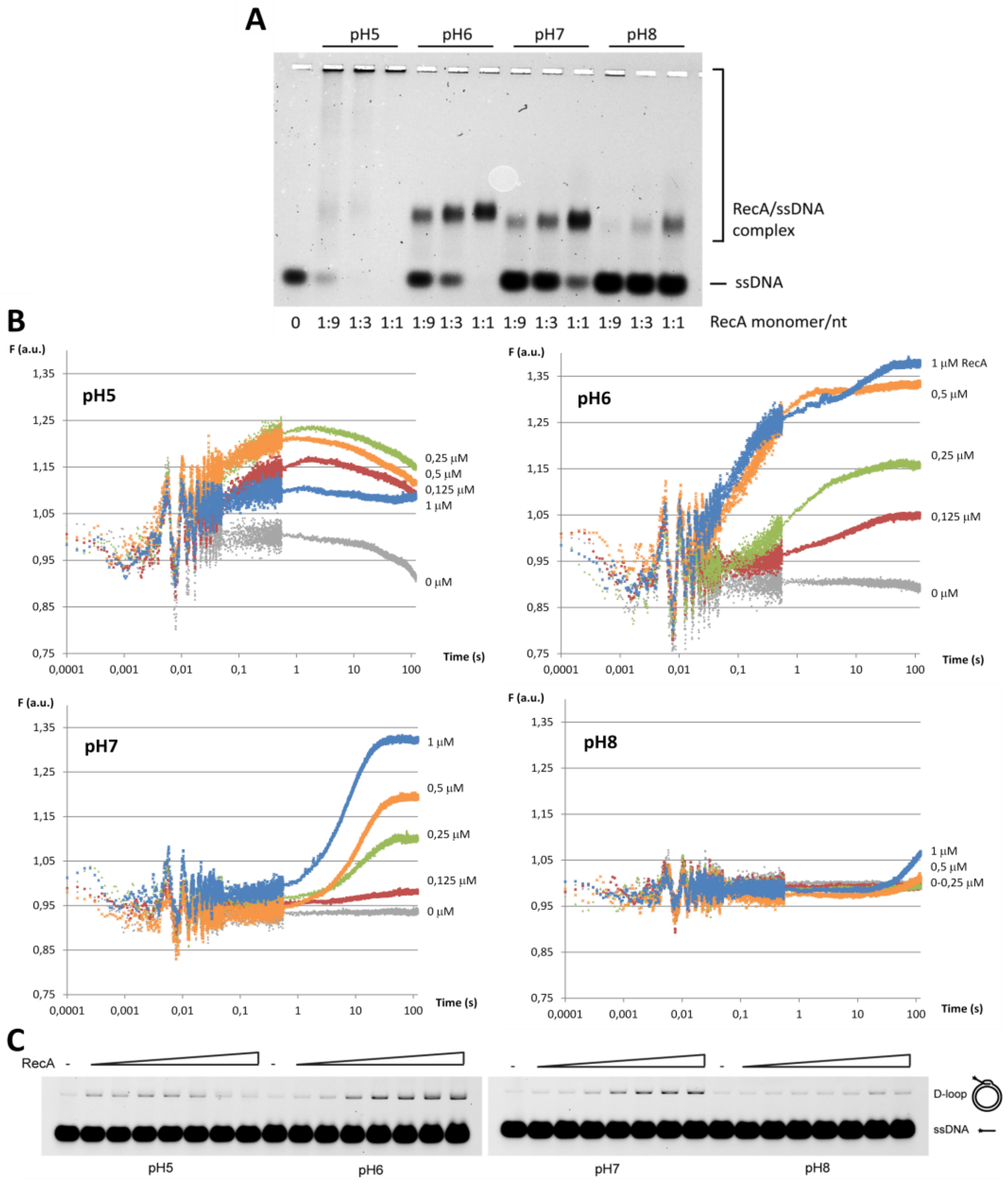
A
pH5
pH6
pH7
pH8
RecA/ssDNA complex
ssDNA
0 1:9 1:3 1:1 1:9 1:3 1:1 1:9 1:3 1:1 1:9 1:3 1:1
RecA monomer/nt
B
F (a.u.)
pH5
0,25 µM
0,5 µM
0,125 µM
1 µM
0 µM
Time (s)
pH6
1 µM RecA
0,5 µM
0,25 µM
0,125 µM
0 µM
pH7
1 µM
0,5 µM
0,25 µM
0,125 µM
0 µM
pH8
1 µM
0,5 µM
0-0,25 µM
C
RecA
D-loop
ssDNA
pH5
pH6
pH7
pH8

**Figure S6. The effect of pH on binding of ssDNA and recombinase activity**

(**A**) Gel illustrating the effect of pH on RecA DNA binding activity. Purified RecA protein was incubated with 90mer FITC-labelled oligonucleotide (10 nM) in the presence of 1 mM ATP and 10 mM $MgCl_2$ for 5 minutes at RT, followed by addition of glutaraldehyde to final concentration 0.125%. Protein-DNA complexes were resolved on 1% agarose gel. RecA monomer versus DNA nucleotide ratio is indicated. Enumeration of the band intensities are shown in Figure 2A.

(**B**) Indicated amounts of RecA protein preincubated with ATP were rapidly mixed with 20 nM 5´Cy3-labelled dT79 oligonucleotide in the presence of 1 mM ATP and 10 mM $MgCl_2$ in stop-flow set-up at various pH conditions. Change of fluorescence was observed from time 0 to 120 sec.

(**C**) Gels illustrating the effect of pH on RecA-mediated D-loop formation. Increasing amounts of RecA protein (0.4, 0.7, 1, 1.5, 2.3, 3.4 and 5.1 μM) were pre-mixed with 50 nM FITC labelled 90mer ssDNA followed by addition of dsDNA plasmid to yield D-loop structure. All samples were then treated with SDS and proteinase K (*SDS/PK*) to release the DNA substrates and analysed by PAGE.

**Figure S7.**

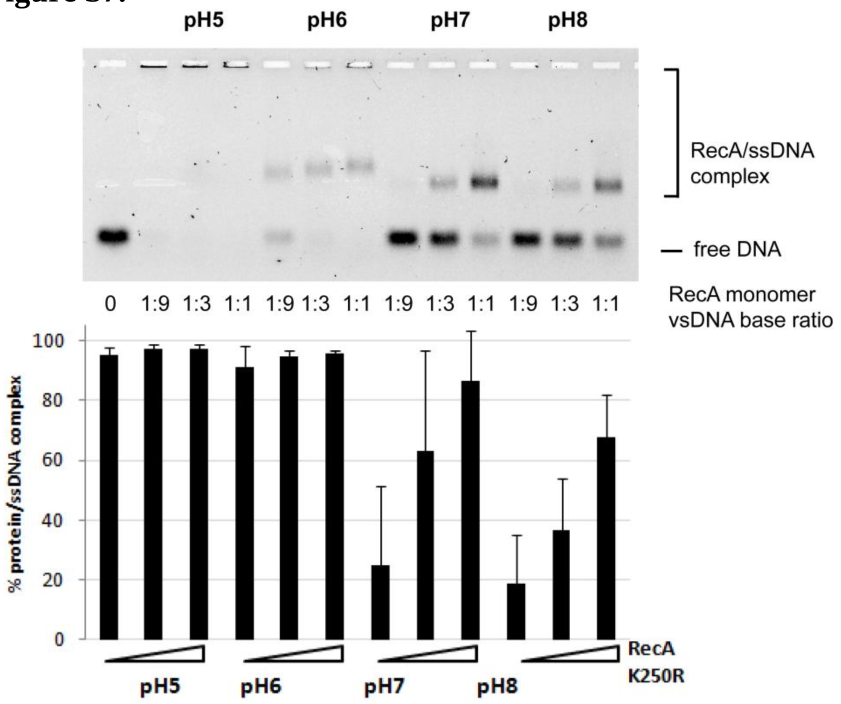
pH5
pH6
pH7
pH8
RecA/ssDNA complex
free DNA
0 1:9 1:3 1:1 1:9 1:3 1:1 1:9 1:3 1:1 1:9 1:3 1:1
RecA monomer vsDNA base ratio
% protein/ssDNA complex
100
80
60
40
20
0
RecA K250R
pH5
pH6
pH7
pH8

**Figure S7. pH dependence of RecA K250R-ssDNA binding.**

Gel illustrating the effect of pH on RecA K250R DNA binding activity. Purified RecA K250R protein was incubated with 90mer FITC-labelled oligonucleotide (10 nM) in the presence of 1 mM ATP and 10 mM $MgCl_2$ for 5 minutes at RT, followed by addition of glutaraldehyde to final concentration 0.125%. Protein-DNA complexes were resolved on 1% agarose gel. RecA monomer versus DNA nucleotide ratio is indicated (mean ± s.d. error bars, ($n$ = 3)).

**Figure S8.**

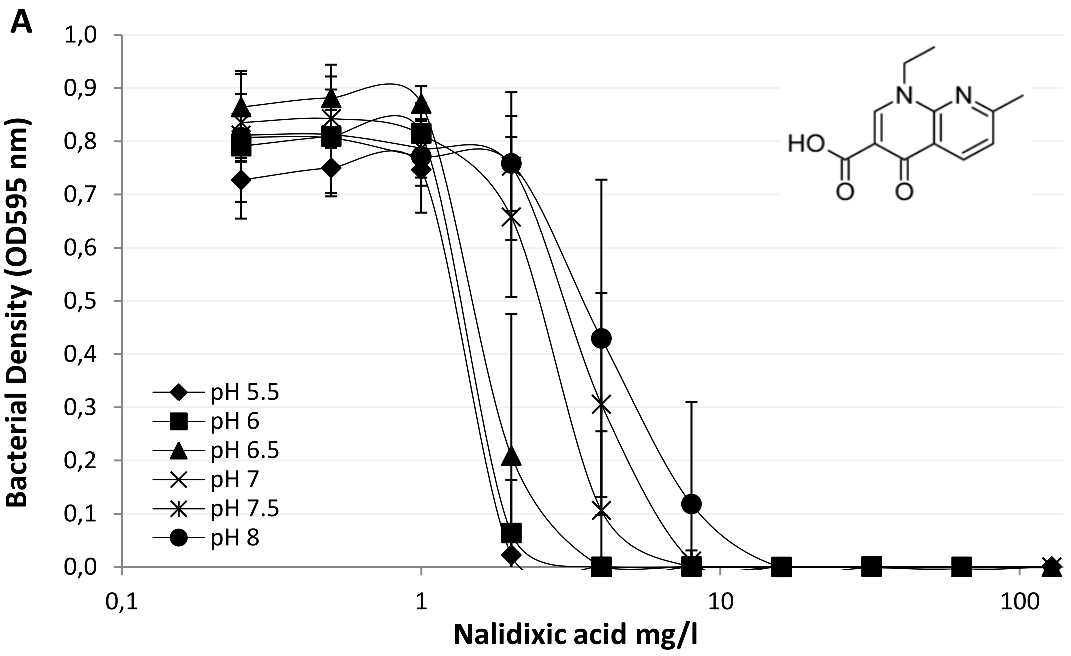
A
1,0
0,9
0,8
0,7
0,6
0,5
0,4
0,3
0,2
0,1
0,0
Bacterial Density (OD595 nm)
pH 5.5
pH 6
pH 6.5
pH 7
pH 7.5
pH 8
0,1
1
10
100
Nalidixic acid mg/l
HO
N
N
O
O

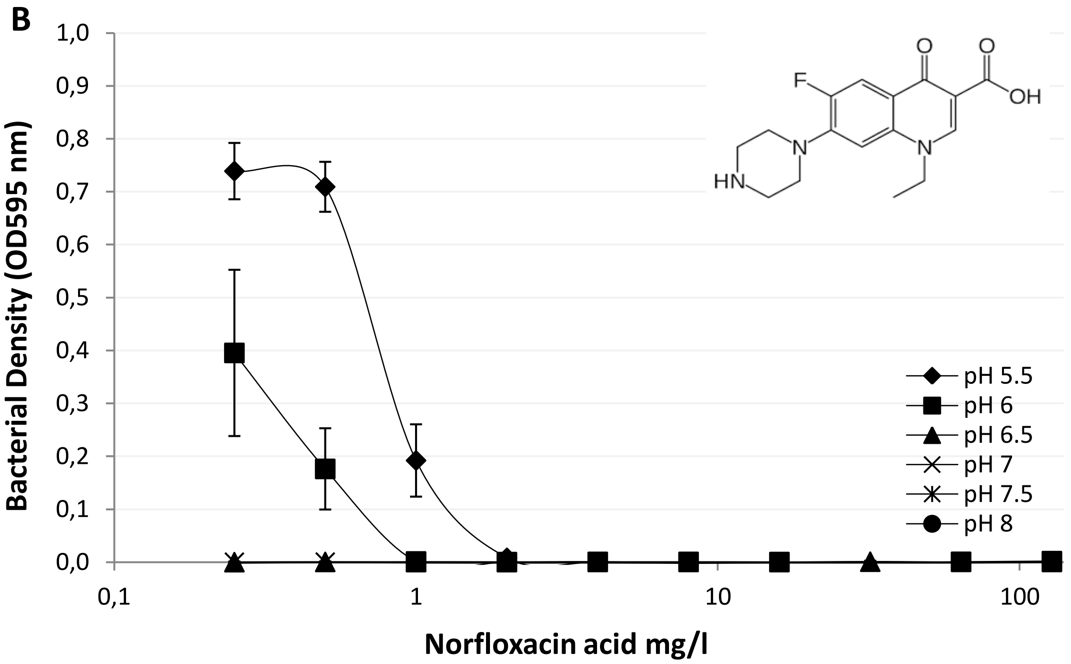
B
1,0
0,9
0,8
0,7
0,6
0,5
0,4
0,3
0,2
0,1
0,0
Bacterial Density (OD595 nm)
F
O
O
OH
N
N
HN
pH 5.5
pH 6
pH 6.5
pH 7
pH 7.5
pH 8
0,1
1
10
100
Norfloxacin acid mg/l

C

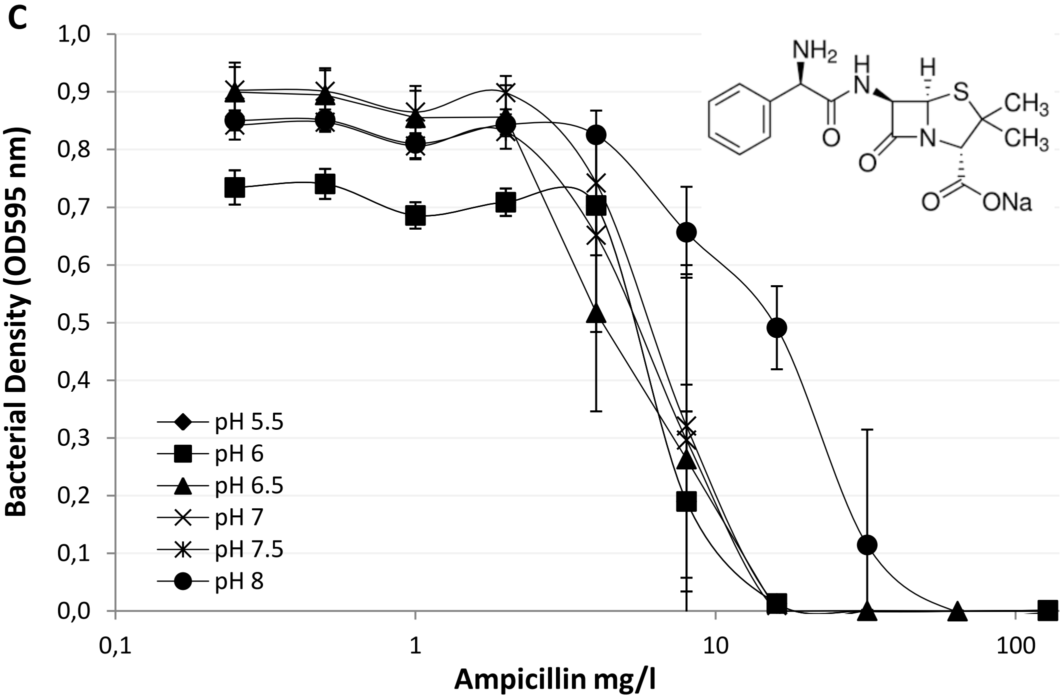

1,0
0,9
0,8
0,7
0,6
0,5
0,4
0,3
0,2
0,1
0,0
Bacterial Density (OD595 nm)
0,1
1
10
100
Ampicillin mg/l
pH 5.5
pH 6
pH 6.5
pH 7
pH 7.5
pH 8
NH2
H
N
H
S
CH3
CH3
O
O
N
O
ONa


D

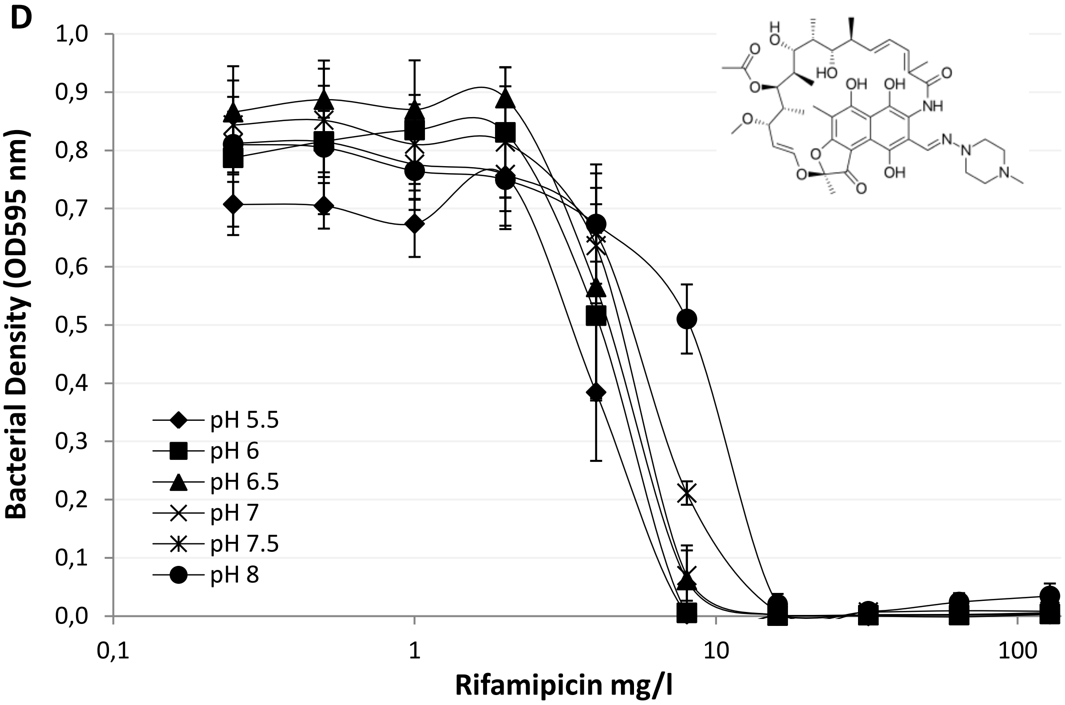

1,0
0,9
0,8
0,7
0,6
0,5
0,4
0,3
0,2
0,1
0,0
Bacterial Density (OD595 nm)
0,1
1
10
100
Rifamipicin mg/l
pH 5.5
pH 6
pH 6.5
pH 7
pH 7.5
pH 8

**E**

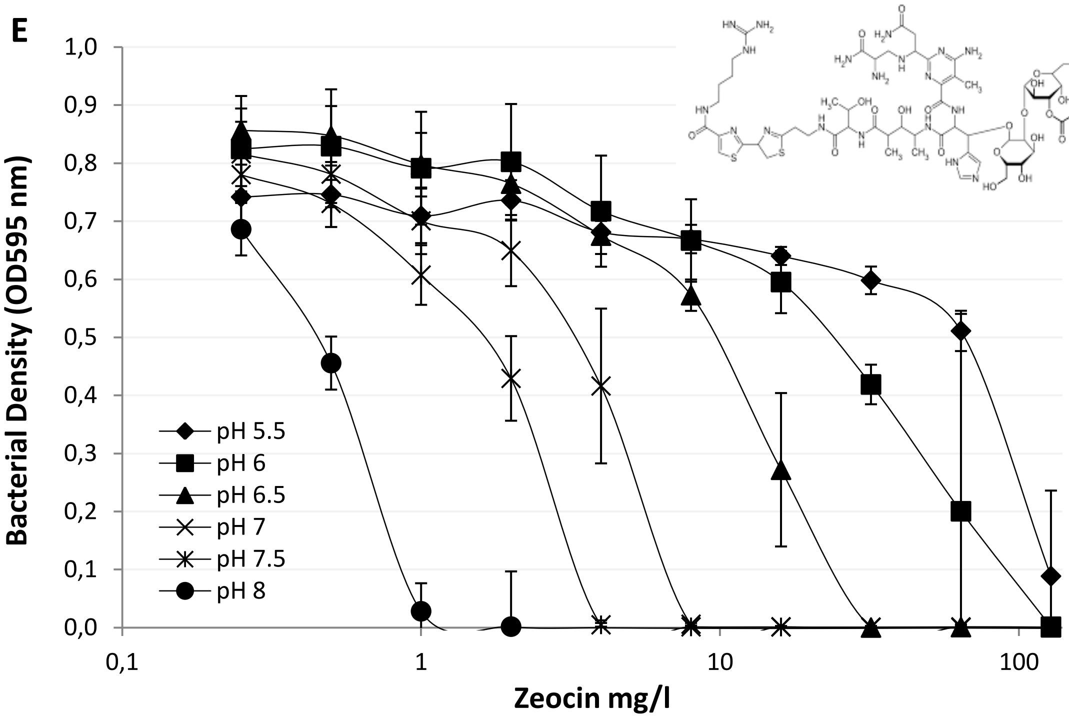

Bacterial Density (OD595 nm)
1,0
0,9
0,8
0,7
0,6
0,5
0,4
0,3
0,2
0,1
0,0
0,1
1
10
100
Zeocin mg/l
pH 5.5
pH 6
pH 6.5
pH 7
pH 7.5
pH 8


**F**

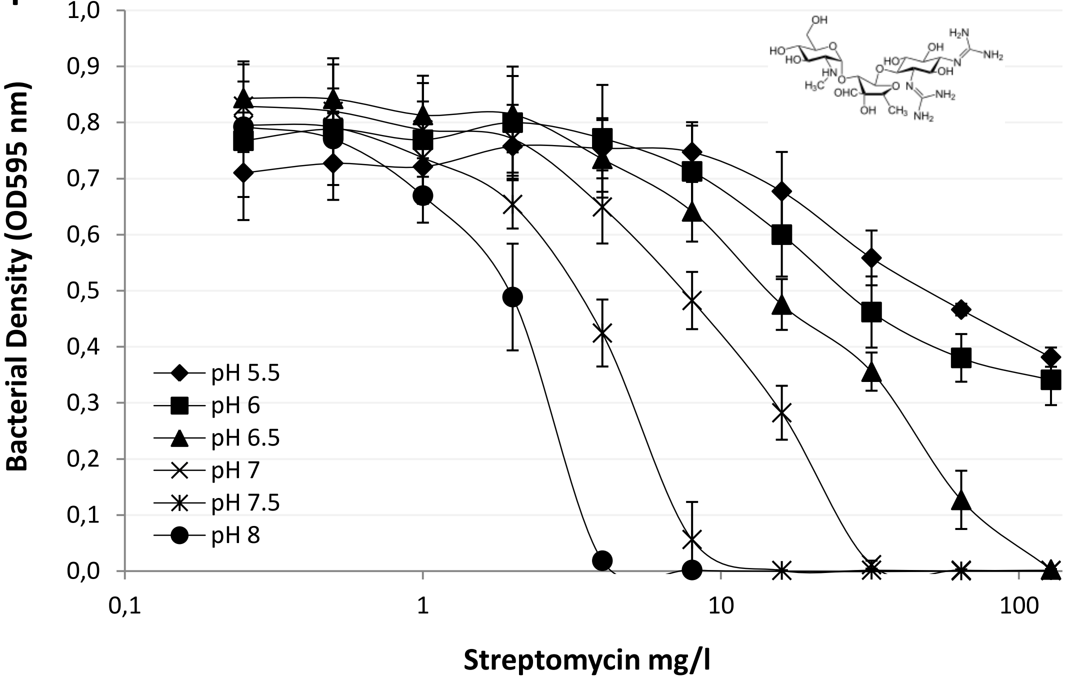

Bacterial Density (OD595 nm)
1,0
0,9
0,8
0,7
0,6
0,5
0,4
0,3
0,2
0,1
0,0
0,1
1
10
100
Streptomycin mg/l
pH 5.5
pH 6
pH 6.5
pH 7
pH 7.5
pH 8

G

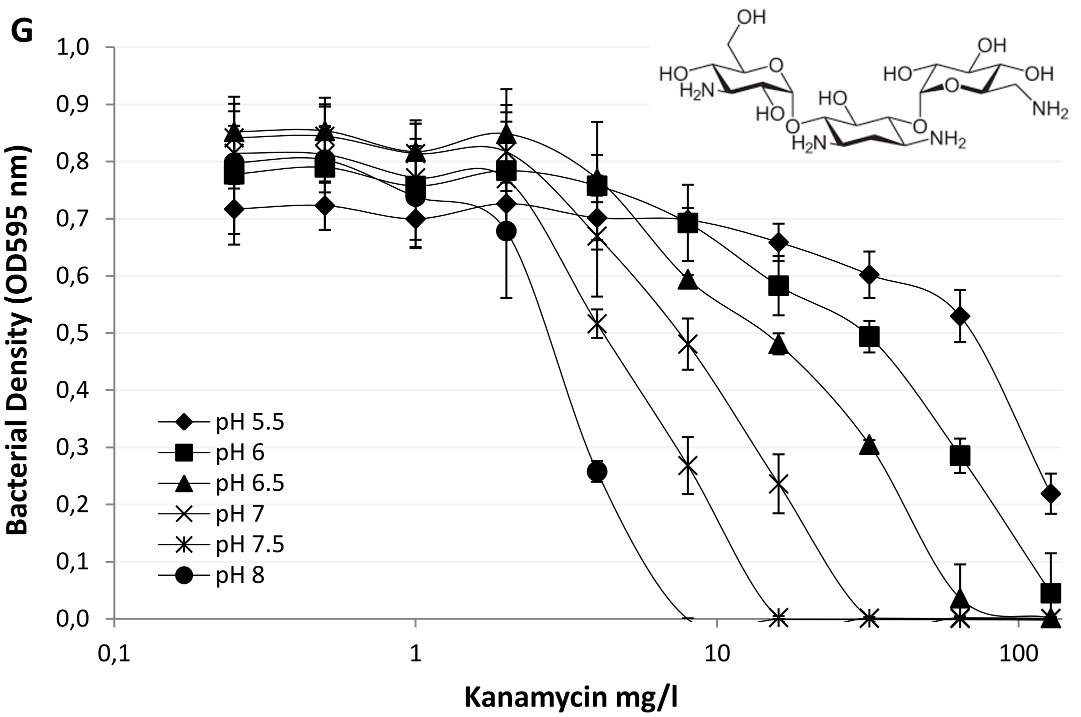

1,0
0,9
0,8
0,7
0,6
0,5
0,4
0,3
0,2
0,1
0,0
Bacterial Density (OD595 nm)
0,1
1
10
100
Kanamycin mg/l
pH 5.5
pH 6
pH 6.5
pH 7
pH 7.5
pH 8


H

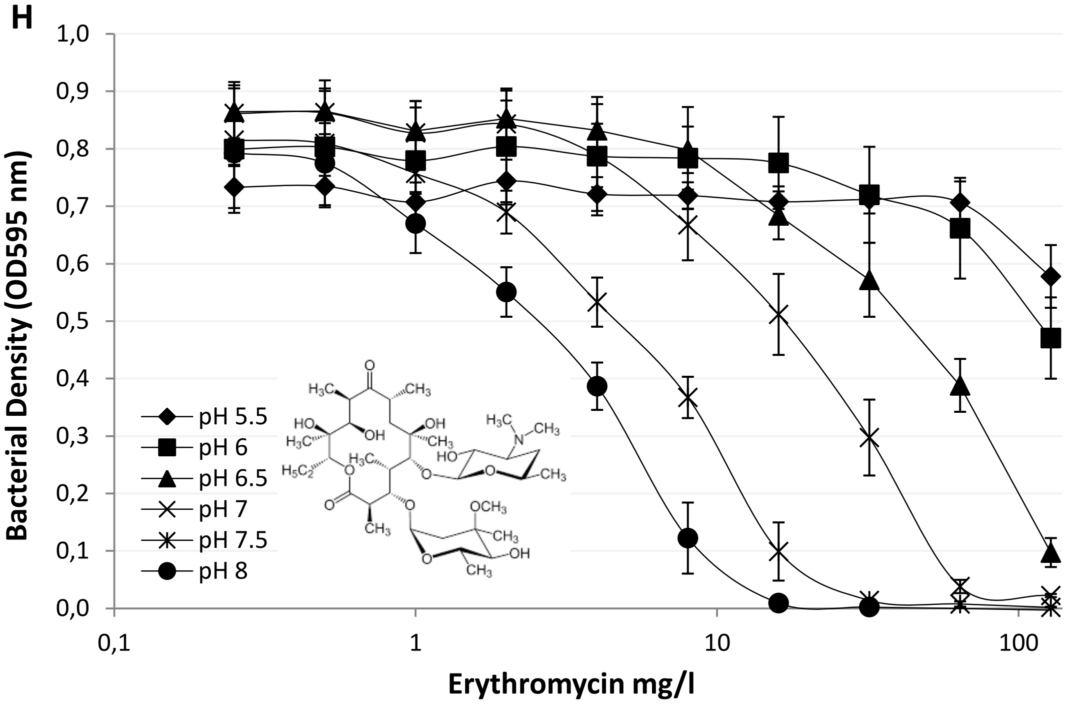

1,0
0,9
0,8
0,7
0,6
0,5
0,4
0,3
0,2
0,1
0,0
Bacterial Density (OD595 nm)
0,1
1
10
100
Erythromycin mg/l
pH 5.5
pH 6
pH 6.5
pH 7
pH 7.5
pH 8

**Figure S8. External pH dependence of the MIC of various classes of antibiotics**

A seed culture of *E. coli* was added to media containing various concentrations of antibiotics (**A**. nalidixic acid, quinolone, **B**. norfloxacin, fluoroquinolone, **C**. ampicillin, beta-lactam, **D**. rifampicin, polyketide/ansamycin, **E**. zeocin, glycopeptides/phleomycin, **F** & **G**. streptomycin and kanamycin respectively, aminoglycosides, **H**, erythromycin, macrolide. After overnight incubation the bacterial density was determined by optical density measurements ($OD_{595}$). All antibiotics showed varying potency over the range of pH's used, however, the range of potency was highest for those containing sugar moieties.

**Figure S9**

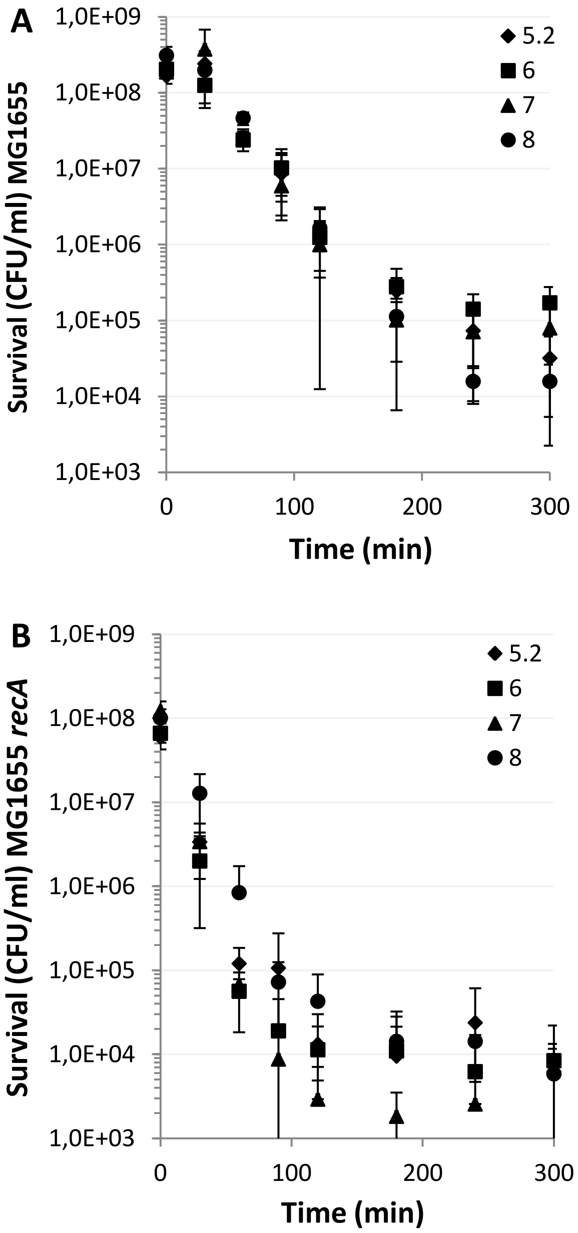
A
1,0E+09
1,0E+08
1,0E+07
1,0E+06
1,0E+05
1,0E+04
1,0E+03
0
100
200
300
Time (min)
Survival (CFU/ml) MG1655
5.2
6
7
8
B
1,0E+09
1,0E+08
1,0E+07
1,0E+06
1,0E+05
1,0E+04
1,0E+03
0
100
200
300
Time (min)
Survival (CFU/ml) MG1655 recA
5.2
6
7
8

**Figure S9. The inherent variation in colony based survival assays made the significance of external pH hard to determine.**

Colony based survival assay of exponentially growth cultures of MG1655 (**A**) and MG1655 *recA* (**B**) following exposure to nalidixic acid 100 µg/ml at time point 0 min at the pH's indicated. Samples removed at the time points indicated were subsequently plated on LB to determine viable colony counts. Data points are mean±s.e.m error bars (n=3 independent replicates).
